# A bird’s white-eye view on neosex chromosome evolution

**DOI:** 10.1101/505610

**Authors:** Thibault Leroy, Yoann Anselmetti, Marie-Ka Tilak, Sèverine Bérard, Laura Csukonyi, Maëva Gabrielli, Céline Scornavacca, Borja Milá, Christophe Thébaud, Benoit Nabholz

**Author notes:** **Cite as:** Leroy T, Anselmetti A, Tilak MK, Bérard S, Csukonyi L, Gabrielli M, Scornavacca C, Milá B, Thébaud C and Nabholz B. A bird’s white-eye view on neo-sex chromosome evolution. bioRxiv 505610, ver. 4 peer-reviewed and recommended by PCI Evolutionary Biology (2019). DOI: 10.1101/505610.

## Abstract

Chromosomal organization is relatively stable among avian species, especially with regards to sex chromosomes. Members of the large Sylvioidea clade however have a pair of neo-sex chromosomes which is unique to this clade and originate from a parallel translocation of a region of the ancestral 4A chromosome on both W and Z chromosomes. Here, we took advantage of this unusual event to study the early stages of sex chromosome evolution. To do so, we sequenced a female (ZW) of two Sylvioidea species, a *Zosterops borbonicus and a Z. pallidus*. Then, we organized the *Z. borbonicus* scaffolds along chromosomes and annotated genes. Molecular phylogenetic dating under various methods and calibration sets confidently confirmed the recent diversification of the genus *Zosterops* (1-3.5 million years ago), thus representing one of the most exceptional rates of diversification among vertebrates. We then combined genomic coverage comparisons of five males and seven females, and homology with the zebra finch genome (*Taeniopygia guttata*) to identify sex chromosome scaffolds, as well as the candidate chromosome breakpoints for the two translocation events. We observed reduced levels of within-species diversity in both translocated regions and, as expected, even more so on the neoW chromosome. In order to compare the rates of molecular evolution in genomic regions of the autosomal-to-sex transitions, we then estimated the ratios of non-synonymous to synonymous polymorphisms (*π*_*N*_*/π*_*S*_) and substitutions (d_*N*_/d_*S*_). Based on both ratios, no or little contrast between autosomal and Z genes was observed, thus representing a very different outcome than the higher ratios observed at the neoW genes. In addition, we report significant changes in base composition content for translocated regions on the W and Z chromosomes and a large accumulation of transposable elements (TE) on the newly W region. Our results revealed contrasted signals of molecular evolution changes associated to these autosome-to-sex chromosome transitions, with congruent signals of a W chromosome degeneration yet a surprisingly weak support for a fast-Z effect.

## Introduction

Spontaneous autosomal rearrangements are common across higher metazoan lineages (Choghlan et al. 2005). By contrast, sex chromosome architectures are much more conserved, even across distant lineages (Murphy et al. 1999; Raudsepp et al. 2004; Fraïsse et al. 2017). A growing number of studies have recently observed departures from this pattern of evolutionary conservation by detecting changes in the genomic architecture of sex chromosomes in some particular lineages – so-called neo-sex chromosomes – that are mainly generated by fusion or translocation events of at least one sex chromosome with an autosome (*e.g.* Kitano & Peichel, 2012; Zhou & Bachtrog, 2012). Considering the long-term conservation of sex chromosome synteny, neo-sex chromosomes provide opportunities to investigate the processes at work during the early stages of sex chromosome evolution (Charlesworth et al. 2005). Previous detailed studies have for example investigated its important role for divergence and speciation (*e.g.* Kitano et al. 2009; Weingartner & Delph, 2014; Yoshida et al. 2014; Bracewell et al. 2017).

From a molecular evolution point of view, an important consequence of the transition from autosomal to sex chromosome is the reduction in effective population size (*N*_*e*_). Assuming a 1:1 sex ratio, *N*_*e*_ of sex-linked regions on the Y (or W) and X (or Z) chromosomes are expected to decrease by three-fourths and one-fourths, respectively (see Ellegren, 2009 for details). According to the neutral theory, nucleotide diversity is then expected to be reduced proportionally to the reduction in *N*_*e*_ due to increased drift effects (Vicoso & Charlesworth, 2006; Pool & Nielsen, 2007). *N*_*e*_ reduction also induces a change in the balance between selection and drift, with drift playing a greater role after the translocation, thus reducing the efficacy of natural selection to purge deleterious mutations from populations. Mutations - including deleterious ones - may also drift to fixation at a faster rate in sex-chromosomes than in autosomes, thus generating expectations for faster evolution at X and Y chromosomes (so-called fast-X or fast-Y effects) (Mank et al. 2007; Rousselle et al. 2016). In addition, given that mutations are on average recessive, positive and negative selection are expected to be more efficient in the heterogametic sex (Hvilsom et al. 2012; Nam et al. 2015). Suppression of recombination is also expected to initiate a degenerative process on the Y chromosome, that may result in the accumulation of nonsynonymous deleterious substitutions owing to a series of processes acting simultaneously: Muller’s ratchet, the Hill-Robertson effect, and linked selection (Charlesworth and Charlesworth, 2000). For the same reason, transposable elements (TEs) are also expected to accumulate soon after the cessation of recombination in the Y chromosomes (Charlesworth 1991; Charlesworth et al. 1994).

Except for few reported examples (de Oliveira et al. 2005; Nanda et al. 2006; Kapusta & Suh, 2017; O’Connor et al. 2018), most birds share a high degree of synteny conservation across autosomal chromosomes (Griffin et al. 2007; Nanda et al. 2008; Ellegren, 2010; Völker et al. 2010; Warren et al. 2010; Ellegren, 2013) and an even higher one at the Z chromosome (Nanda et al. 2008). A notable exception is the neo-sex chromosome of Sylvioidea species, a superfamily of passerine birds in which a translocation of a large part of the Zebra finch 4A chromosome onto both the W and the Z chromosomes, as characterized by genetic mapping (hereafter translocations of the neoW-4A and neoZ-4A on ancestral W and Z sex chromosomes; Pala et al. 2012a). Terminologically, these original sex chromosomes (W or Z) are considered as specific regions of the neoW and neoZ chromosomes (hereafter neoW-W and neoZ-Z). All along the manuscript, we have used this terminology to emphasize the fact that these translocations also induce substantial evolutionary shifts on original sex chromosomes. These two autosome-to-sex chromosome transitions are present in reed warblers (Acrocephalidae), old-world warblers (Sylviidae) and larks (Alaudidae) (see also Brooke et al. 2010), and therefore likely occurred in the common ancestor of all present-day Sylvioidea (Pala et al. 2012a,b; Sigeman et al. 2018), between 15 and 30 million years ago (Myrs) (Ericson et al. 2014; Prum et al. 2015; Nabholz et al. 2016). These two sex chromosome translocations provide unique opportunities to investigate the early stages of the W and Z chromosome evolution.

Based on phylogenetic trees calibrated using geological events (Moyle et al. 2009), *Zosterops* species of the family Zosteropidae (more commonly referred to as white-eyes) are considered to have emerged around the Miocene/Pliocene boundary. Considering both this recent emergence and the remarkable high diversity currently observed in this genus (more than 80 species), this group appears to have one of the highest diversification rates reported to date for vertebrates and is therefore considered as one of the ‘great speciator’ examples (Diamond et al. 1976; Moyle et al. 2009). White-eye species are typical examples of taxa spanning the entire “grey zone” of speciation (Roux et al. 2016). As a consequence of these different degrees of reproductive isolation between taxa, white-eyes have long been used as models to study bird speciation (*e.g.* Clegg and Philimore 2010; Melo et al. 2011; Oatley et al. 2012; Oatley et al. 2017). Among all white-eyes species, the Reunion grey white-eye *Zosterops borbonicus* received considerable attention over the last 50 years. This species is endemic from the volcanic island of Reunion and shows an interesting pattern of microgeographical variation, with five distinct colour variants distributed over four specific regions across the 2,500 km^2^ of island surface. Both plumage color differentiation data (*e.g.* Gill et al. 1973; Milá et al. 2010; Cornuault et al. 2015) and genetic data (*e.g.* Milá et al. 2010; Delahaie et al. 2017) support this extensive within-island diversification. Despite its important role in the understanding of the diversification of *Zosterops* species, no genome sequence is currently available for this species. More broadly, only one *Zosterops* species has been sequenced to date (the silvereye *Z. lateralis,* Cornetti et al. 2015).

Here, we obtained detailed genome data from *Z. borbonicus*, a member of the Zosteropidae family and then arranged scaffolds into pseudochromosomes to provide insights into the evolutionary processes that may have contributed to the early stages of the sex chromosome evolution. We also generated a more fragmented genome sequence for *Z. pallidus* for molecular dating and molecular evolution analyses. We found similar breakpoint locations for both translocated regions suggesting evolution from the same initial gene sets, and studied the molecular evolution of the two newly sex-associated regions. By comparing levels of within-species nucleotide diversity at autosomal and sex chromosomes, we found support for a substantial loss of diversity on both translocated regions, largely consistent with expectations under neutral theory. We then compared patterns of polymorphisms and divergence at neo-sex chromosome genes and found support for a considerable fast-W effect, but surprisingly weak support for a fast-Z effect. Investigations of candidate changes in base composition led to the identification of specific signatures associated with abrupt changes in recombination rates (reduction or cessation) of the two neo-sex regions. Finally, we reported higher transposable elements (TE) content on the newly W than on the newly Z regions, suggesting ongoing neoW chromosome degeneration.

## Results

### *Zosterops borbonicus* Genome Assembly

Using a strategy combining long-read sequencing with PacBio and short-read Illumina sequencing with both mate-pairs and paired-end reads, we generated a high-quality reference genome for a female Reunion grey white-eye captured during a field trip to Reunion (Mascarene archipelago, southwestern Indian Ocean). The 1.22 gigabase genome sequence comprises 97,503 scaffolds (only 3,757 scaffolds after excluding scaffolds smaller than 10 kbp), with a scaffold N50 of 1.76 Mb (Fig. 1, Table S1). The completeness of the assembly is very high based on the BUSCO statistic (93.0%). Among all investigated avian species, the ‘GRCg6a’ chicken genome assembly is the only one exceeding this value (93.3%) (Fig. S1). Compared to the other species, our *Z. borbonicus* reference assembly also exhibits the lowest proportions of ‘missing’ (2.5%, a value only observed for the reference chicken genome) and ‘fragmented’ genes (4.3%, a value which is 0.1% higher than for the reference chicken assembly).

**Figure 1:**
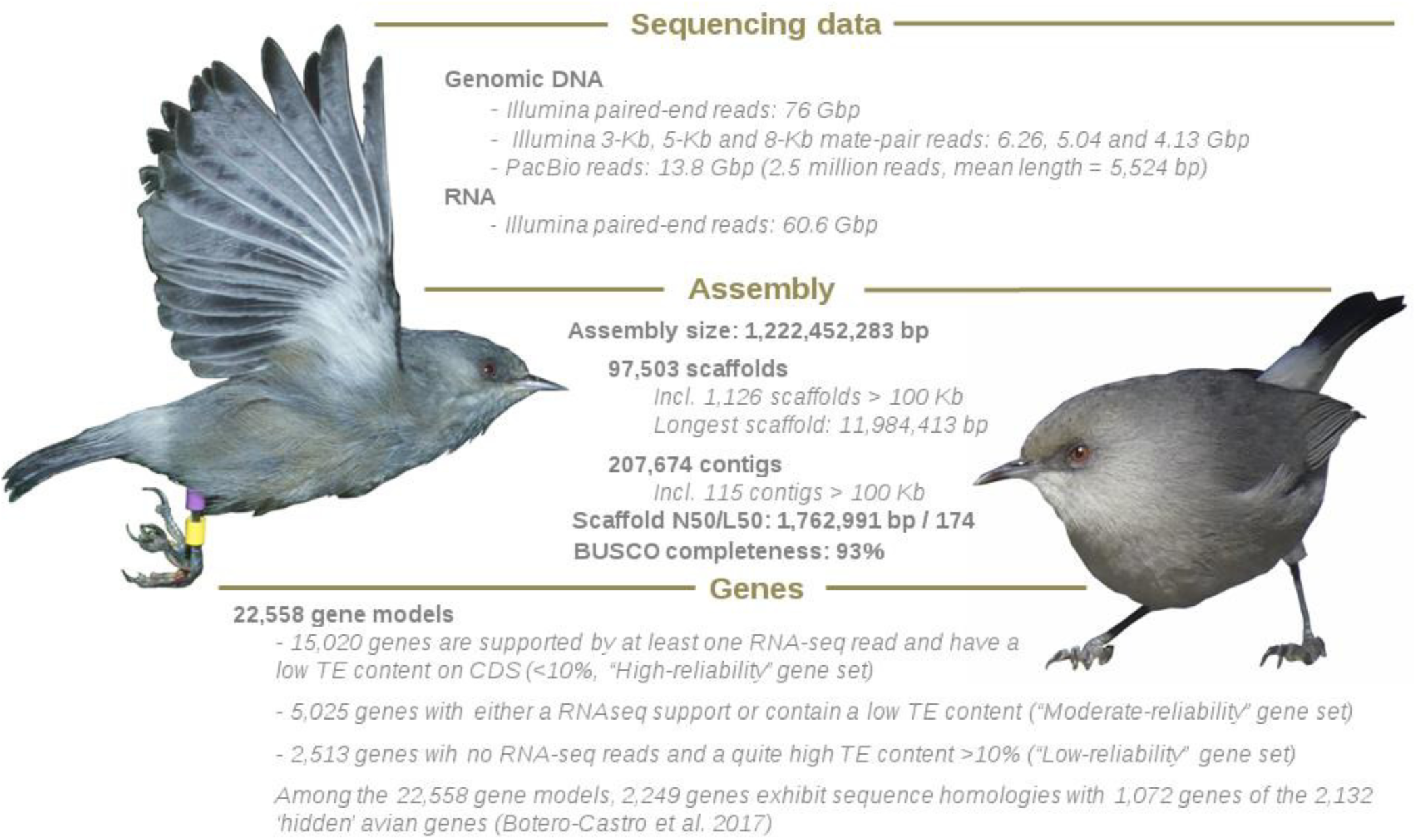
*Z. borbonicus* genome assembly and gene model summary statistics. The reported numbers of reads and the total numbers of bases sequenced correspond to the trimmed reads. The three different gene sets are indicated as specified in the gff file. Photo credits: Maëva Gabrielli & Laurent Brillard (http://laneb.re)

Using the PASA pipeline combined with EVidenceModeler (Haas et al. 2008; Haas et al. 2011) and several *in silico* tools trained by the RNAseq data, a total of 22,558 gene models were predicted. Among these 22,558 genes, half of the “hidden” avian genes identified by Botero-Castro et al. (2017), *i.e.* a large fraction of avian genes in GC-rich regions missing from most avian genome assemblies, were recovered (1,072 out of 2,132).

The vast majority of the 22,558 CDS (83.3%) are supported by at least one RNA-seq read (18,793 CDS), including 17,474 with a FPKM above 0.01. Among the 22,558 gene models, TE content is globally low (8.5%), but 4,786 genes exhibit a TE content in coding regions greater than 0.25, including 1,512 genes predicted to be TE over the total length of the CDS (Fig. S2). Still more broadly, the level of expression is strongly negatively correlated with the within-gene TE content (r^2^=0.038, p<2.2×10^−16^, after excluding 709 genes with FPKM>100).

Additionally, we generated a genome assembly of a female Orange River white-eye (*Z. pallidus*) using 10X mate-pair and 72X paired-end reads after cleaning. This assembly is much more fragmented (170,557 scaffolds and scaffold N50 = 375 Kb, Table S1) than the *Z. borbonicus*. Both *Z. borbonicus* and *Z. pallidus* assemblies are available from Figshare repository URL: https://figshare.com/s/122efbec2e3632188674 (see also the data availability section).

### Reference-assisted genome organization

We then anchored scaffolds using the v3.2.4 reference genome of the zebra finch (Warren et al. 2010) assuming synteny. We used the zebra finch as a pivotal reference since this reference sequence is of high-quality, with 1.0 of 1.2 gigabases physically assigned to 33 chromosomes including the Z chromosome, plus three additional linkage groups based on genetic linkage and BAC fingerprint maps (Warren et al. 2010). We anchored 928 among the longest scaffolds to the zebra finch chromosomal-scale sequences, thus representing a total of 1.01 Gb among the 1.22 Gb of the *Z. borbonicus* assembly (82.8%).

In parallel to the zebra finch-oriented approach, we used DeCoSTAR (Duchemin et al. 2017), a tool that improves the assembly of several fragmented genomes by proposing evolutionary-induced adjacencies between scaffolding fragments containing genes. To perform this analysis, we used the reference sequences of 27 different avian species (Table S2) with associated gene tree phylogeny of 7,596 single copy orthologs (trees are available from the following Figshare repository, URL: https://figshare.com/s/122efbec2e3632188674). Among the 97,503 scaffolds (800 containing at least one orthologous gene), DeCoSTAR organized 653 scaffolds into 188 super-scaffolds for a total of 0.837 Gb (68.5% of the *Z. borbonicus* assembly), thus representing a 2.59-fold improvement of the scaffold N50 statistic (4.56 Mb). Interestingly, among the 465 scaffold junctions, 212 are not only supported by gene adjacencies within the other species, but are also supported by at least one *Z. borbonicus* paired-end read. From a more global point of view, DeCoSTAR not only improved the *Z. borbonicus* genome, but also those of 25 other species (mean gain in scaffold N50 over 11 Mb, representing 3.30-fold improvement on average, range: 1.0-6.02). The only exception is the already well-assembled chicken genome reference. For all these species, the proposed genome organizations (“agp files”) were made available at the following URL: https://figshare.com/s/122efbec2e3632188674.

We then combined zebra finch-oriented and DeCoSTAR approaches for the *Z. borbonicus* genome, by guiding DeCoSTAR using the a priori information of the zebra finch-oriented approach to get beyond two limitations. First, DeCoSTAR is a gene-oriented strategy, and thus cannot anchor scaffolds without genes that have orthologous analogues in the other species, which is generally the case for short scaffolds. Second, the zebra-finch oriented approach assumes a perfect synteny and collinearity between *T. guttata* and *Z. borbonicus*, which is unlikely. By combining both approaches, we were able to anchor 1,082 scaffolds, including 1,045 scaffolds assigned to chromosomes representing a total 1.047 Gb (85.7% of the *Z. borbonicus* assembly). In addition, DeCoSTAR helped propose more reliable *Z. borbonicus* chromosomal organizations for these 1,045 scaffolds by excluding some *T. guttata*-specific intra-chromosomal inversions.

### Assigning scaffolds to W and Z Chromosomes

To identify sex chromosome scaffolds, we first mapped trimmed reads from males and females *Z. borbonicus* individuals which were previously sequenced by Bourgeois et al. (2017) and then computed median per-site coverage over each scaffold for males and females (Fig. 2). After taking into account differences in coverage between males and females, we then identified scaffolds that significantly deviated from 1:1 and identified neoW and neoZ scaffolds (see methods section). This strategy led to the identification of 218 neoW (7.1 Mb) and 360 neoZ scaffolds (91.8 Mb) among the 3,443 scaffolds longer than 10 kb (Fig. 2). Among the 360 neoZ scaffolds assigned by coverage, 339 scaffolds were already anchored to the neoZ chromosome based on the synteny-oriented approach, thus confirming the accuracy of our previous assignation and suggesting that we have generated a nearly complete Z chromosome sequence The list of scaffolds identified on the two neo-sex chromosomes was made available: https://figshare.com/s/5ad54809ed89dba83db7.

**Figure 2:**
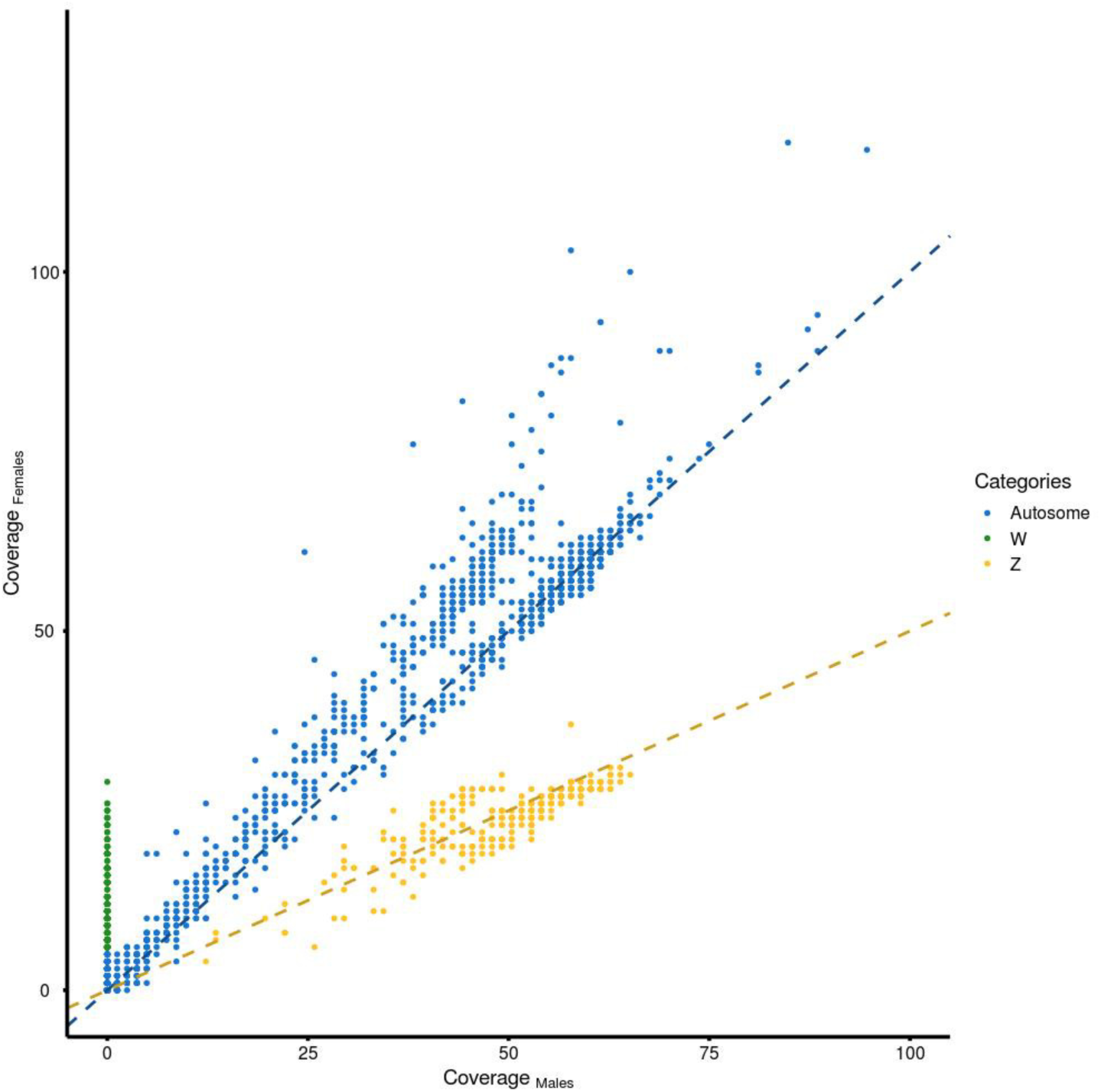
Identification of Z and W scaffolds using coverages of male and female individuals. Normalized sum of median per-site coverage of trimmed reads over the five males (x-axis) and seven females (y-axis). Values were only calculated for scaffolds longer than 10-Kb. Linear regressions for slope=1 (blue) and slope=0.5 (yellow) are shown. Scaffolds were assigned following the decision rule described in the material and methods section.

Due to the absence of a chromosomal-scale W sequence for a passerine bird species, we are unable to provide a chromosomal structure for the 218 neoW scaffolds. We however assigned 174 among the 218 neoW scaffolds to the neoW-4A region. These scaffolds representing a total length of 5.48 Mb exhibited high levels of homology with the zebra finch 4A chromosome. We found support for neoW-4A scaffolds aligning between positions 21,992 and 9,603,625 of the *T. guttata* 4A chromosome (thus representing 57% of the corresponding 9.6 Mb 4A region). Leaving aside the difficulty of sequencing and assembling the neoW chromosome (Tomaszkiewicz et al. 2017), a large part of the difference between the total assembled size and the corresponding region in *T. guttata* is likely due to a 1.75 Mb chromosomal deletion on the neoW-4A. Indeed, we found no neoW scaffold with a homology to the large 4A *T. guttata* region between positions 1,756,080 and 3,509,582. Based on the *T. guttata* reference genome, this region was initially gene-poor, since only two *T. guttata* genes were found in this large genomic region (*i.e.* 1.1 genes/Mb), as compared to the 142 genes observed on the whole translocated region (*i.e.* 14.8 genes/Mb). Remaining neoW scaffolds, *i.e.* those having no homology with the 4A *T. guttata* region, were considered as sequences belonging to the ancestral W chromosome (hereafter neoW-W region), except for six scaffolds showing reliable hits but only at specific locations of the scaffold, and for which the accuracy of any assignation was considered too low.

### Molecular dating

The molecular phylogenetic analyses were aimed at estimating the divergence time of the *Zosterops* genus. Indeed, even if our focal dataset is composed of only three species, the divergence between *Z. pallidus / Z. borbonicus* and *Z. lateralis* represents the first split within the genus except *Z. wallacei, i.e.* the origin of clade B in Moyle et al. (2009). As a consequence, our phylogeny (Fig. 3, S3) is expected to provide an accurate estimate for the onset of the diversification of the *Zosterops* lineage. For this molecular dating, we added three species to the 27 species used in the previous analyses and generated gene sequence alignments. Given that these newly added species - namely the silvereye, Orange River white-eye and willow warbler - have no gene models available yet, we used the AGILE pipeline (Hughes & Teeling 2018, see method) to obtain orthologous sequences. Due to the inherent computational burden of Bayesian molecular dating analyses, we randomly selected 100 alignments among the least GC rich single-copy orthologs and performed ten replicated analyses (chronograms available at FigShare URL: https://figshare.com/s/122efbec2e3632188674). Indeed, GC poor genes are known to be slowly and more clock-like evolving genes as compared to the other genes (Jarvis et al. 2014).

**Figure 3:**
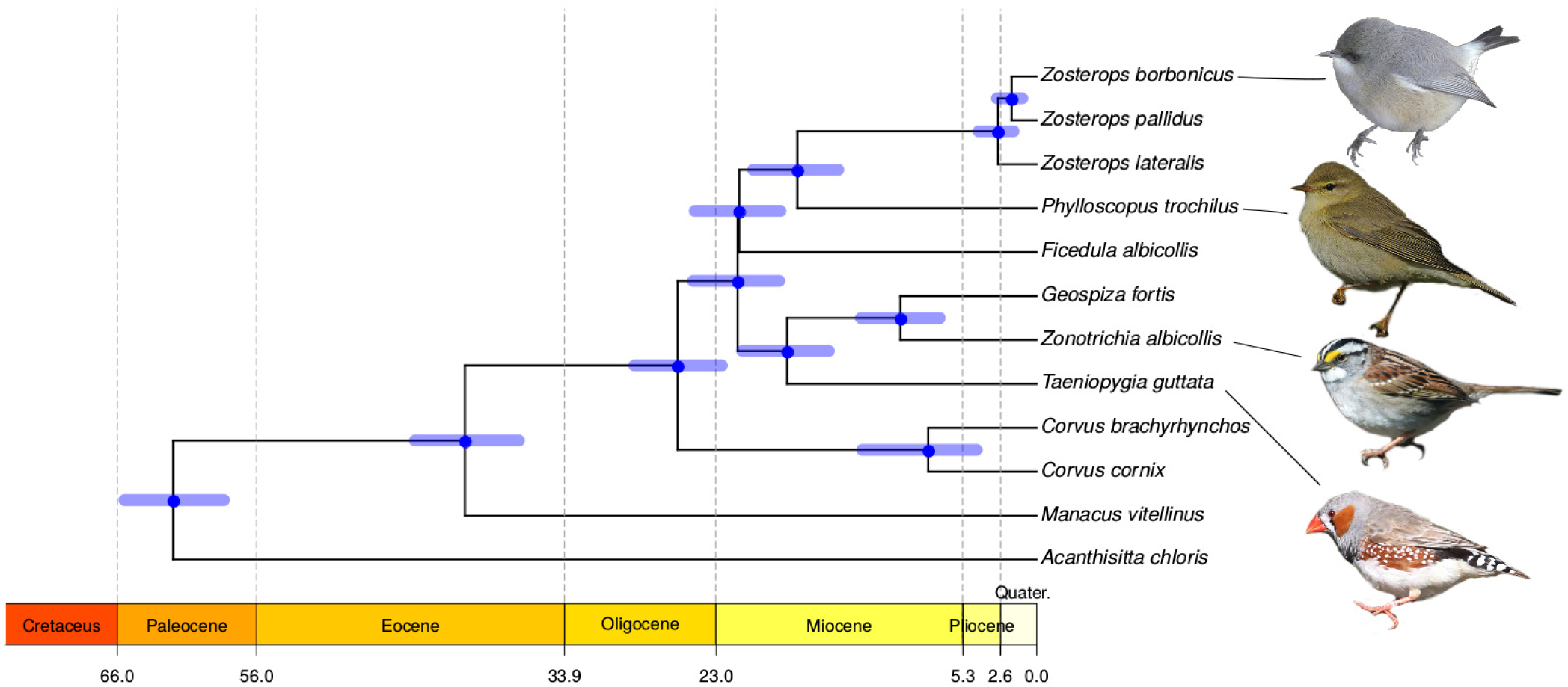
Molecular phylogeny and dating of the investigated passerine birds. Estimates of molecular dating are based on dataset 1 and calibration set 1 (Table S3). For greater clarity, the cladogram focuses on passerine birds only, but see Fig S3 for a cladogram of the inferred phylogeny for the whole set of investigated avian species.

We used several combinations of fossil calibrations and substitution models (Table S3). For the radiation of Neoaves, all analyses led to molecular dating consistent with Jarvis et al. (2014) and Prum et al. (2015) with estimates around 67-70 Myrs, except for the calibration set 4 (82 Myrs), albeit with large 95% confidence intervals (CI = 64 - 115 Myrs) (Table 1). Calibration set 4 is very conservative with no maximum calibration bound except for the Paleognathae / Neognathae set to 140 Myrs. In contrast, calibration set 3 is the more constrained with the Suboscines / Oscines split bounded between 28 - 34 Myrs. Unsurprisingly, this calibration led to the youngest estimates, dating the origins of passerines at 59 Myrs (56 - 63 Myrs) and the Paleognathae / Neognathae at 65 Myrs (63 - 69 Myrs).

**Table 1:**
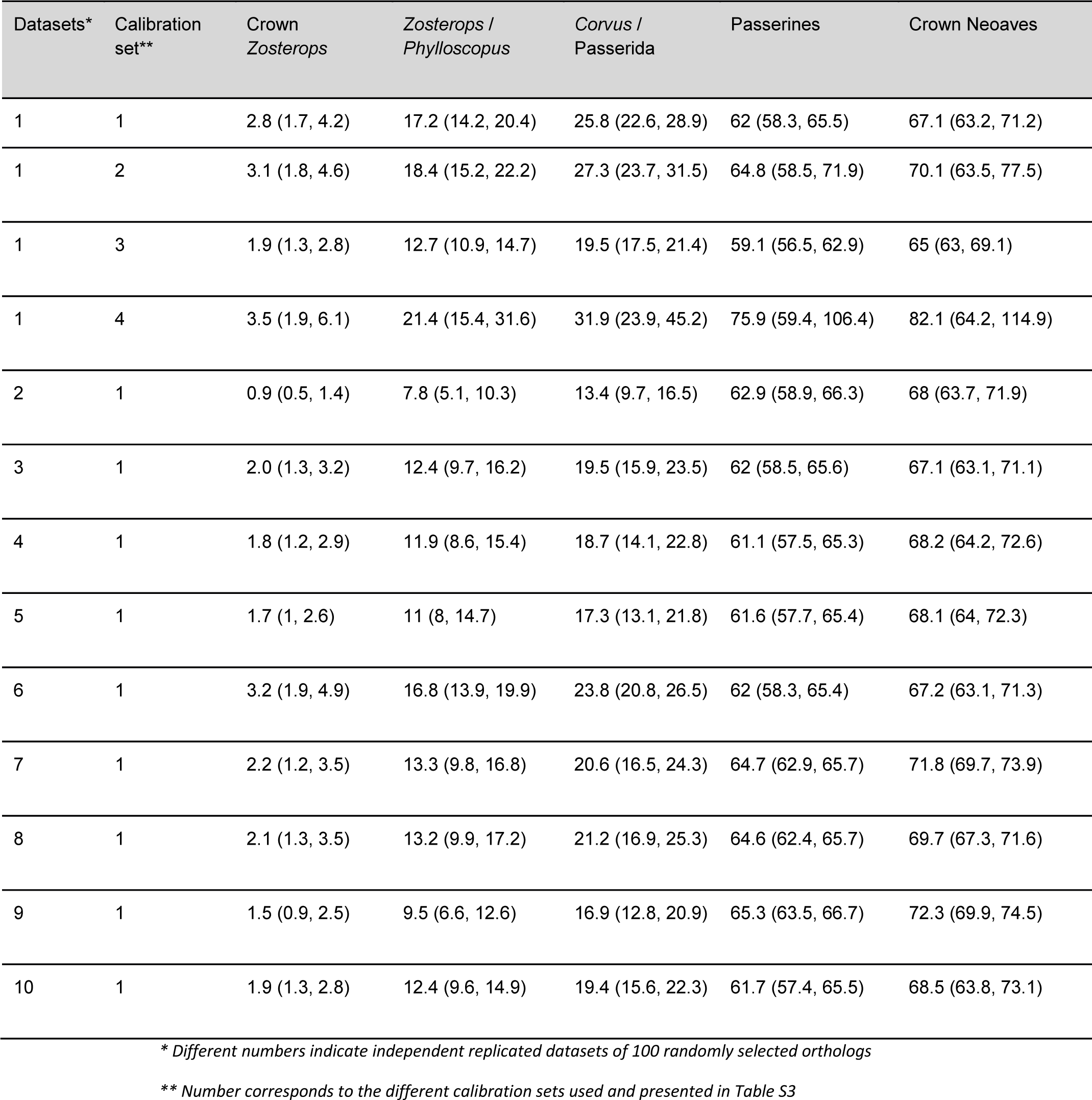
Molecular dating analyses. Mean dates are indicated and 95% confidence intervals (CI) are provided within parentheses

In all runs, our estimates of the origin of *Zosterops* have lower limits of CI including 2.5 Myrs and mean estimates are also often considerably lower than this value (Table 1). Using the calibration set 1, the ten replicated datasets gave a mean age estimate for the origin of *Zosterops* around 2 million years ago (Table 1). More broadly, our analysis is consistent with a recent origin of *Zosterops*, with a crown clade age of less than 5 Myrs and probably between 1 and 3.5 Myrs (Table 1).

Additionally, we performed a completely independent molecular dating by applying the regression method proposed in Nabholz et al. (2016). Based on nine full *Zosterops* mitochondrial genomes, we calculated molecular divergence. The divergence between *Z. lateralis* and the other *Zosterops* has a median of 0.205 subst./site (max = 0.220, min = 0.186) for the third codon position. Assuming a median body mass for the genus of 10.7 g (Dunning, 2007), we estimated a divergence date between 2.3 and 6.2 Myrs. These estimates are in line with our previous dating based on nuclear data and with molecular dating based on fossil calibration confirming the extremely rapid diversification rate of white-eyes.

### Chromosomal breakpoints

Our synteny-oriented approach using pairwise whole-genome alignments of *Z. borbonicus* and *T. guttata* sequences helped us in identifying the chromosomal breakpoint of the neoZ sex chromosome. We reported a scaffold (scaffold329) with long sequence alignments with both the 4A and the Z chromosomes (Fig. 4). Considering the intervals between the last LASTZ hit on the 4A and the first one on the Z chromosome, we estimated that this breakpoint occurred between positions 9,605,374 and 9,606,437 of the 4A zebra finch chromosome (genome version: v.3.2.4), which is fairly close to the estimate of 10 Mb previously reported by Pala et al. (2012a). Based on the soft-masked version 3.2.4 of the zebra finch genome, this 1-Kb region is well assembled (no ambiguous “N” bases) and shows no peculiarities in TE or GC content (7.4% and 39.0%, respectively) as compared to the rest of the zebra finch 4A chromosome (18.7% and 43.7%).

**Figure 4:**
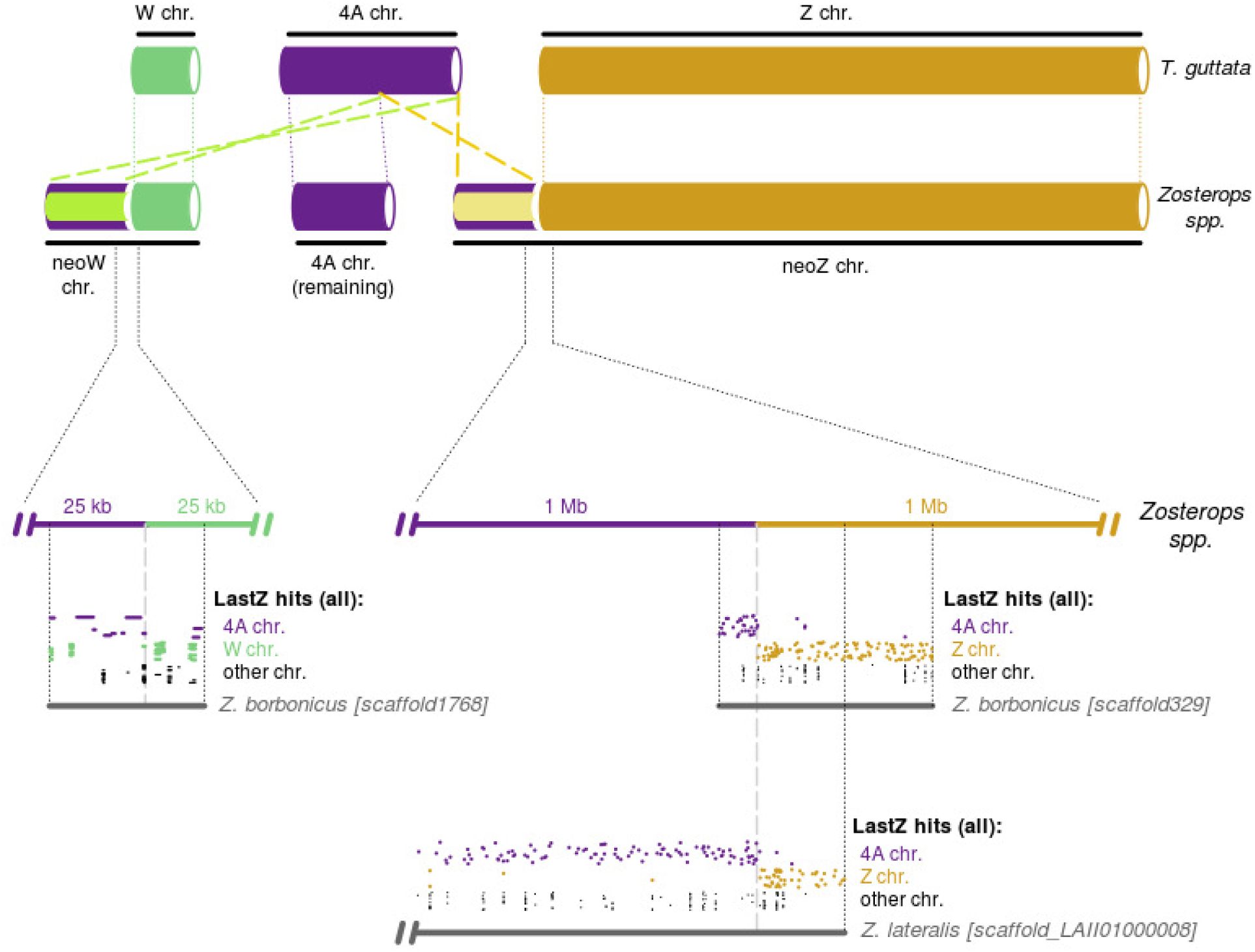
Fine-scale genome architecture of the Sylvioidea neo-sex chromosomes. Original W and Z and autosomal 4A chromosomes are shown in green, yellow and purple, respectively. Translocated regions of the ancestral 4A chromosome (neoW-4A and neoZ-4A) are shown with a purple outline. LASTZ hits on scaffold supporting the chromosomal breakpoints are shown using the same color code. For ease of illustration, coordinates on the Z and 4A chromosomes are reversed (i.e. the white disk represents the beginning of the *T. guttata* chromosome sequence).

To discard the potential confounding factor of a sequence artifact due to a chimerism in the *Z. borbonicus* assembly, we used the procedure for the sequence assembly of *Z. lateralis* (Cornetti et al. 2015) and identified a scaffold (LAII01000008.1) supporting the same chromosomal breakpoint. Using the *Z. lateralis* sequence, we estimated that this breakpoint occurred between positions 9,605,524 and 9,606,431.

We then investigated the chromosomal breakpoint for the neoW (Fig. 4). We identified a candidate scaffold, *scaffold1768_size33910*, with alignment hits on both the 4A *T. guttata* and the W of *F. albicollis* (Smeds et al. 2015). Among all W scaffolds, this scaffold aligns at the latest positions of the 4A *T. guttata* chromosome (from 9,590,067 to 9,603,625), which is remarkably close to our estimation for the region translocated on the neoZ chromosome. Considering also alignments against contigs of the *F. albicollis* W chromosome, we estimated that the chromosomal breakpoint probably occurred between positions 9,603,625 and 9,605,621.

### Chromosomal-scale estimates of nucleotide diversity

We used sequencing data from six *Z. borbonicus* individuals (three males and three females) sequenced by Bourgeois et al. (2017) to explore the genomic landscape of within-species diversity. Over all autosomal 10-Kb windows, mean nucleotide diversity (Tajima’s π; Tajima 1983) estimates were roughly similar in males and females (π_males_=1.82e-3 and π_females_=1.81e-3, respectively). The nucleotide diversity landscape greatly varies within and between chromosomes (A in Fig 5, Fig S4 & S5). In addition, we identified some series of 10-Kb windows with very low level of nucleotide diversity (red bars, Fig. 5A). Interestingly, some of these regions also exhibit the highest negative values of Tajima’s D (red bars, Fig. 5B & S6; *e.g.* the end of the chromosome 2). Small interchromosomal differences in the distribution of nucleotide diversity values were detected for both female- and male-based estimates, suggesting that both datasets give similar results at the chromosomal level, except for the neoZ chromosome for which substantial differences were observed between the two datasets (Fig. S4). Even considering this source of variability, the neoZ chromosome still shows significant deviation from the mean autosomal diversity for both datasets (π_females_=1.04e-3 and π_males_=1.34e-3, respectively; t-tests, p<2e-16 for both datasets), thus representing 57.6% and 73.5% of the mean autosomal diversity. This reduced level of nucleotide diversity was similarly detected for the neoZ-Z and the neoZ-4A regions of the neo-Z chromosome (C, Fig. 5). Based on both datasets, a lower nucleotide diversity was observed on neoZ regions as compared to the autosomal chromosomes. Mean π_males_ was estimated to 1.21e-3 for the neoZ-4A region and 1.35e-3 for the neoZ-Z region, corresponding to 66.6% and 74.2% of the autosomal diversity. Mean π_females_ values are roughly similar with 1.37e-3 and 1.00e-3, thus representing 76.2% and 55.5% of the autosomal diversity, respectively.

**Figure 5:**
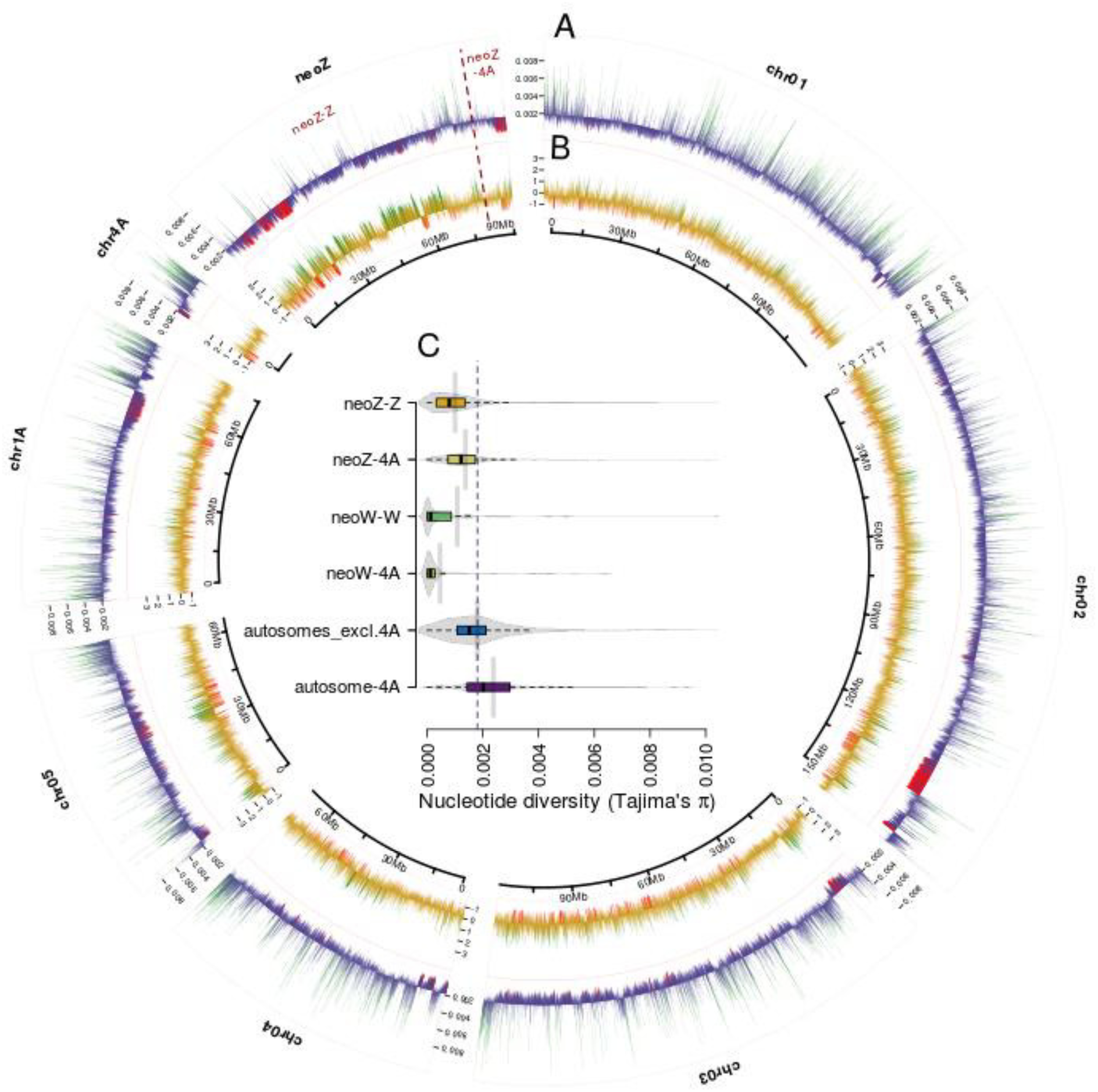
Intra- and inter-chromosomal variations in nucleotide diversity. Variations of Tajima’s π_males_ (A) and D_males_ (B) estimates along the six Z. borbonicus macrochromosomes, the autosomal 4A and the neo-Z chromosome. The two metrics were calculated in non-overlapping 10-Kb sliding windows. Top 2.5% and bottom 2.5% windows are shown in green and red, respectively. For both Tajima’s π_males_ and Tajima’s D_males_, each bar shows the deviation from the mean genomic value over the whole genome (Tajima’s π_males_ and Tajima’s D_males_ baselines: 1.82e-3 and - 0.26, respectively). C) Interchromosomal differences in Tajima’s π_females_ between autosomes and sex chromosomes. Fig. S5 and S6 show Tajima’s pi and D along most Z. borbonicus chromosomes.

Similarly, data from females only were used to estimate the level of within-species diversity variation on the neoW chromosome (C, Fig. 5). We similarly observed a reduced level of diversity as compared to the autosomal chromosomes (mean π_females_=5.87e-4; t-test, p<2e^−16^), with only one-third (32.5%) of the mean nucleotide diversity estimated for the autosomes. Differences in Tajima’s π_females_ were observed on the neoW-W region of the neoW chromosome and on the neoW-4A translocated region, albeit non-significant (t-test, p=0.102), with higher diversity on the neoW-W (π=1.08e-3) as compared to the neoW-4A (π=4.65e-4). Median values are however much more consistent between the two regions (π=1.25e-4 and π=1.16e-4, respectively), suggesting that few neoW-W windows greatly contributed to this discrepancy (C, Fig. 5).

### Ratios of non-synonymous to synonymous polymorphisms (π_n_/π_s_) and substitutions (d_n_/d_s_)

We computed *π*_*N*_*/π*_*S*_ ratios among all genes in autosomal, neoZ and neoW chromosomes. Estimated *π*_*N*_*/π*_*S*_ of chromosome 4A genes is slightly lower as compared to the rest of autosomal chromosomes (*π*_*N*_/*π*_*S*_= 0.212 vs. *π*_*N*_*/π*_*S*_ = 0.170). NeoZ-Z exhibits slightly higher values than both autosomal sets (*π*_*N*_*/π*_*S*_=0.282), however no or very little difference has been observed between neoZ-4A (*π*_*N*_*/π*_*S*_ = 0.181) and autosomal chromosomes. On the contrary, *π*_*N*_*/π*_*S*_ ratios on genes of the neoW chromosome are very high, with *π*_*N*_*/π*_*S*_=0.418 for the neoW-4A and *π*_*N*_*/π*_*S*_= 0.780 for the neoW-W regions.

We estimated *d*_*N*_*/d*_*S*_ ratios for *Z. borbonicus* and ten additional passerine species (all passerines except *A. chloris* in Fig. 1) for a total of 6,339 alignments of single-copy orthologs, corresponding to 6,073 autosomal genes, 66 on the remaining autosomal region of the 4A chromosome (hereafter autosome-4A), 164 on the neoZ-Z, 54 on the neoZ-4A and 17 on the neoW-4A. For the neoZ-4A and neoW-4A, we made a special effort to identify gametologs (homologous sequences between neoZ-4A and neoW-4A genes, which were previously excluded during the filtering of 1:1 orthologs).

We then compared the *d*_*N*_*/d*_*S*_ of neoZ-4A and neoZ-Z genes. To do that, we randomly subsampled the data to match the number of genes in neoZ-4A region (54) before concatenation and then computed *d*_*N*_*/d*_*S*_ ratios. The variability in *d*_*N*_*/d*_*S*_ is evaluated by bootstrapping genes and creating repeated concatenation of 54 genes for each genomic region.

For all species, the *d*_*N*_*/d*_*S*_ ratio reaches significantly higher values for neoZ-Z genes than for autosomal genes (Fig. 6). Surprisingly, we found no evidence for an increase in *d*_*N*_*/d*_*S*_ ratios in the neoZ-4A when compared to autosomes (Fig. 6, species in the red frame). For both the willow warbler and *Zosterops* species, *d*_*N*_*/d*_*S*_ ratios are lower in neoZ-4A genes than in autosomal regions (willow warbler: neoZ-4A *d*_*N*_*/d*_*S*_ = 0.09 (95% CI=0.07-0.11) *vs.* autosomal *d*_*N*_*/d*_*S*_ = 0.11 (95% CI=0.08-0.15); for white-eyes: neoZ-4A *d*_*N*_*/d*_*S*_ = 0.10 (95% CI = 0.08-0.13) *vs.* autosomal *d*_*N*_*/d*_*S*_ = 0.14 (95% CI=0.09-0.20)). This also holds true when we compare genes of the ancestral 4A chromosome translocated on the neoZ chromosome (neoZ-4A, “chromosome 4A:0-9.6 Mb” in Fig. 6) and genes of the 4A chromosome (“chromosome 4A:9.6-20.7 Mb”). Even if we report slightly higher *d*_*N*_*/d*_*S*_ for the translocated region as compared to the rest of the 4A chromosome, such a difference between the two ancestral 4A regions is also observed in several other species that do not have the translocation (e.g., *M. vitellinus, T. gutatta* or *Z. albicollis;* Fig. 6), including a much bigger difference for *T. guttata* (Fig. 6). As a consequence, our d_N_/d_S_ analysis did not provide any support for a higher d_N_/d_S_ ratio associated to the autosomal-to-Z translocation.

**Figure 6:**
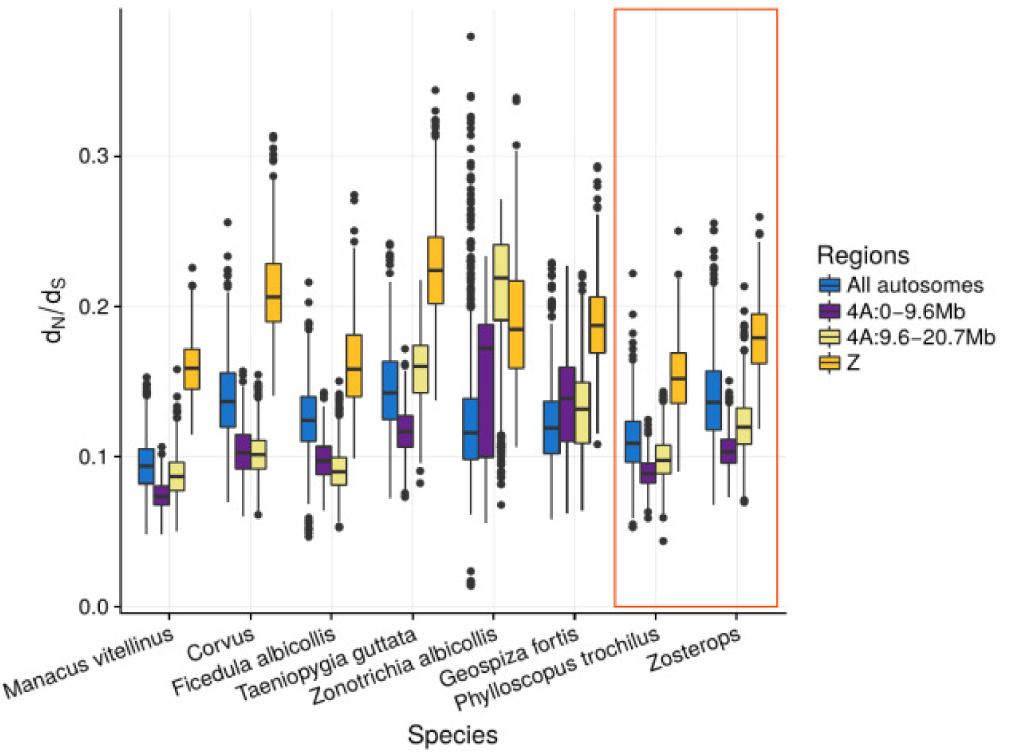
Variation in d_N_/d_S_ ratios among chromosomal regions for the 11 investigated species. Estimates are performed at the genus level, i.e. on branches before the split of the two *Corvus* (*C. cornix* and *C. brachyrhynchos*) and the three *Zosterops* species (*Z. lateralis, Z. pallidus* and *Z. borbonicus*). The red box shows species with the translocated neoZ-4A region For these species, the 4A:0-9.6 Mb region corresponds to the neoZ-4A and the Z corresponds to the neoZ-Z.

Next, we computed the *d*_*N*_*/d*_*S*_ of the branch leading to the neoZ-4A and neoW-4A copies in the *Zosterops borbonicus* genome. In this case, the *d*_*N*_*/d*_*S*_ of the neoW-4A genes were significantly higher than the *d*_*N*_*/d*_*S*_ of the neoZ-4A copies (mean *d*_*N*_*/d*_*S*_ = 0.531, sd = 0.417 for neoW-4A genes; mean *d*_*N*_*/d*_*S*_ = 0.194, sd = 0.234 for neoZ-4A genes; Wilcoxon signed rank test, p-value =7.7e-5, Fig. 7). The increase in *d*_*N*_*/d*_*S*_ is particularly strong, including some neoW-4A genes with *d*_*N*_*/d*_*S*_ close or slightly higher than 1.

**Figure 7:**
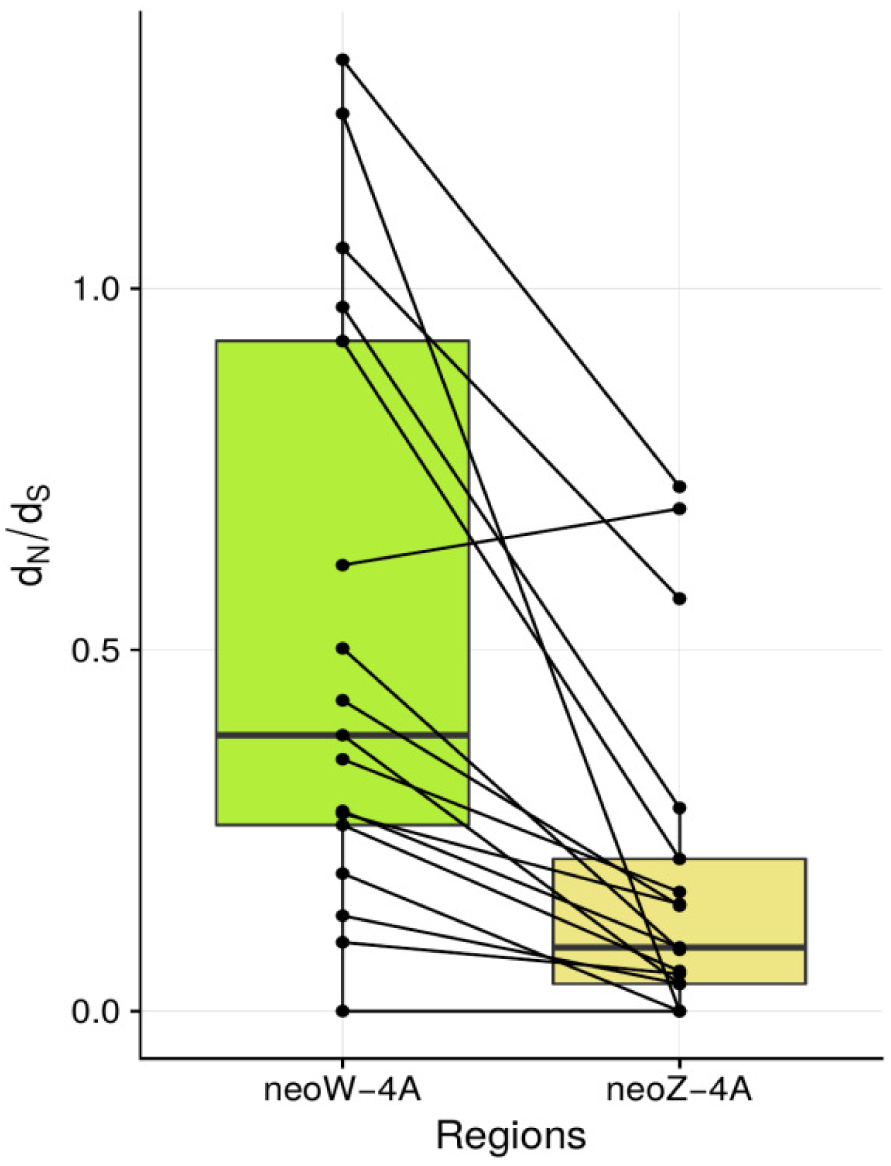
The d_N_/d_S_ ratios of the neoW-4A and neoZ-4A copies (gametologs) of the *Zosterops borbonicus* genomes. For each gene, horizontal lines linked values of the two gametologs.

For the three genes with a *d*_*N*_*/d*_*S*_ >1, we performed a likelihood-ratio test comparing a model with a fixed *d*_*N*_*/d*_*S*_ value equaling 1 (null model) to a model with a dN/dS value free to vary. All observed values were not significantly different from the null model. Based on these tests, these results are therefore more consistent with ongoing pseudogenization than positive selection. However, it should be outlined that we detected no premature stop-codon or frameshift mutation in the six neoW-W genes with the highest dN/dS values (i.e. *d*_*N*_*/d*_*S*_ reaching at least 0.5).

### GC, GC* and Transposable Elements (TE) Contents

Next, we investigated the potential change in base composition after the translocation events, due to changes in recombination rates and more precisely, the recombination-associated effect of GC-biased gene conversion (gBGC, Duret & Galtier 2009, Nabholz et al. 2011, Weber et al. 2014). For this purpose, we computed the GC content over 10-Kb sliding windows (Fig. S7) and the GC content at equilibrium for orthologous sequences (Fig. S8). As expected, differences in GC contents are observed between chromosomes, with a higher GC content in short chromosomes. Among all chromosomes, the neoZ chromosome showed the second lowest median GC rate. Interestingly, GC content is lower in the neoZ-4A regions as compared to the autosomal 4A chromosome (Student’s t-test, p=2.2e-4) or as compared to the neoW-4A, although this difference is only marginally significant (t-test, p=0.059). Slight differences in GC content are observed between neoW-4A (t-test, p=0.066) and neoW-W (t-test, p=0.096) as compared to the autosomal-4A chromosome suggesting that the GC content of the neoW-4A likely decreased after the translocation, but to a lesser extent than for the neoZ-4A (Fig. 8). We also evaluated variation in GC content at equilibrium (GC*) for the new gametologs, *i.e.* homologous genes in the neoW-4A and neoZ-4A regions (Fig. 9A). For these 17 genes, all Passerida species without the neo-sex regions exhibited a GC* between 0.7 and 0.8 (Fig. 9A). Passerida species with these neo-sex chromosomes showed a slightly reduced GC* at neoZ-4A genes (mean GC* = 0.57 and 0.68 for the willow warblers and the white-eyes respectively, and a strongly reduced GC* content at neoW-4A genes (mean=0.4, 95% CI = 0.30-0.51, Fig. 9A). In contrast, the GC* of the non-translocated region of the ancestral chromosome 4A (position 9.6-20.7 Mb) apparently remains unchanged between the Sylvioidea and the other Passerida (GC* between 0.71 and 0.86, Fig. 9B).

**Figure 8:**
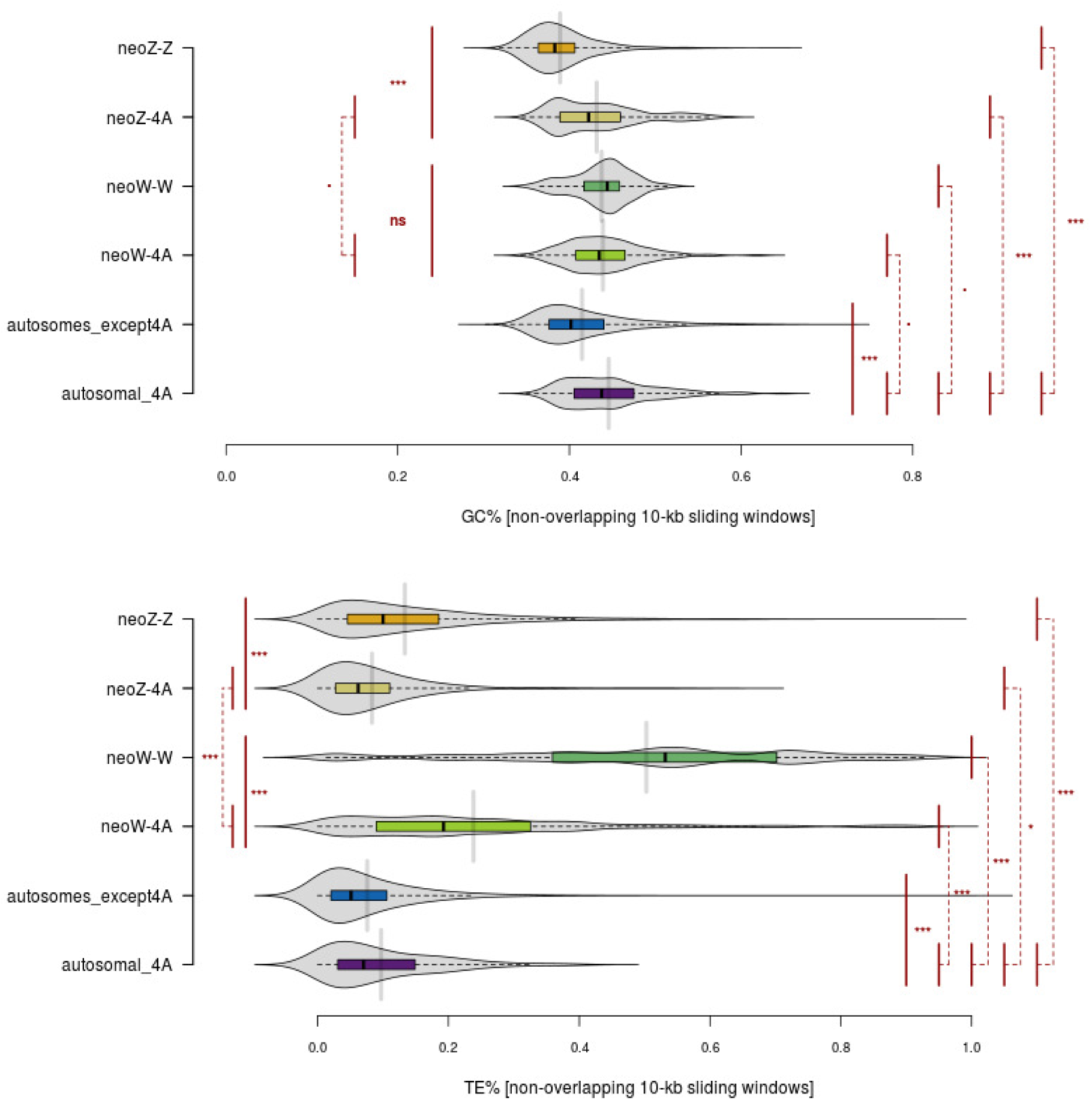
Variation in GC (top) and TE (below) contents between chromosomes. GC and TE contents were estimated along the whole genome using non-overlapping 10-kb sliding windows. Means were compared using t-tests (pairwise comparisons between chromosome sets, without corrections for multiple testing). Thresholds:. < 0.1, * < 0.05, ** < 0.01, *** < 0.001

**Figure 9:**
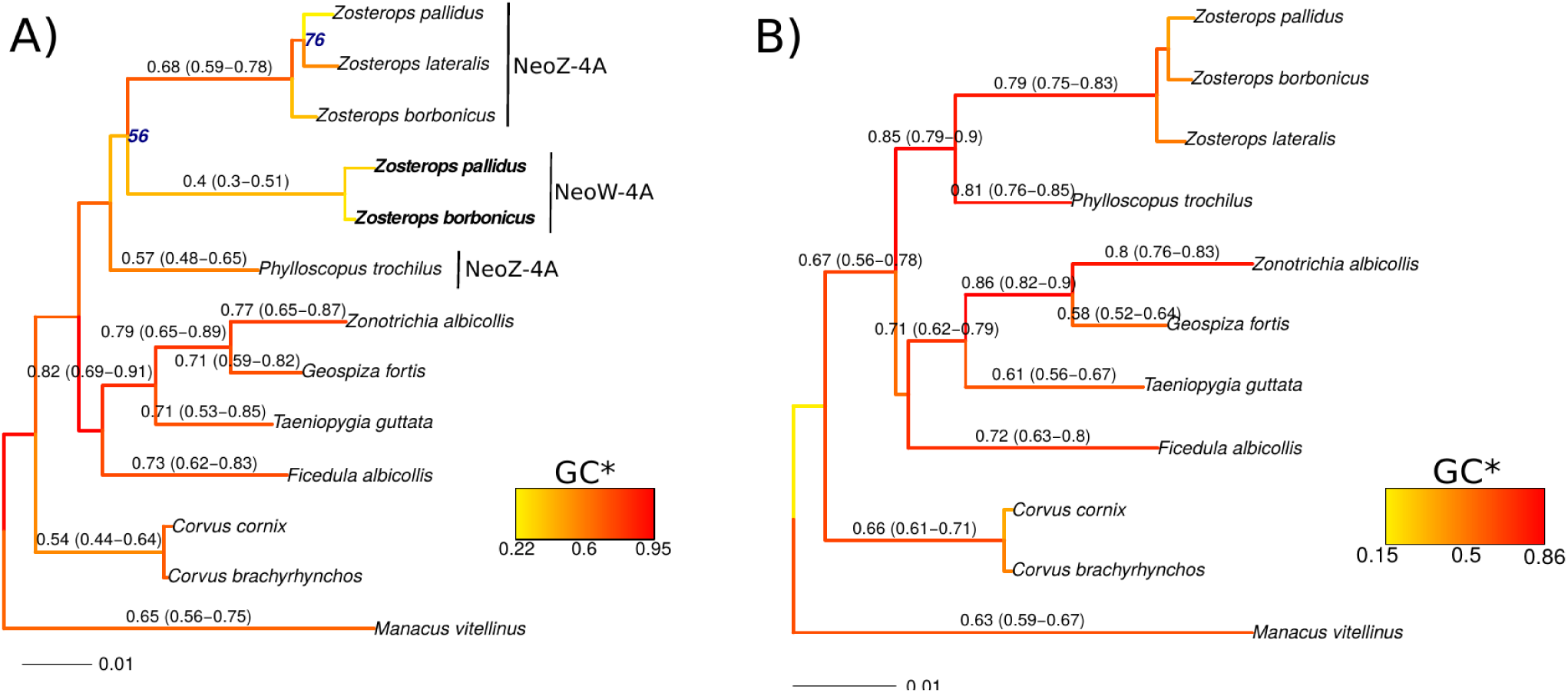
Phylogenetic relationships and estimation of GC equilibrium (GC*). A) Estimation using the 17 genes coded by the translocated region of the ancestral chromosome 4A (0-9.6 Mb). Bold font indicates NeoW-4A sequences. B) Estimation using the 79 genes coded by the non-translocated region of the ancestral chromosome 4A (9.6-20.7 Mb). Branch color and numbers above branches indicated GC*, mean and 95% confidence interval obtained by bootstrap are in parenthesis. Number in blue in front of the node indicates ultra-fast bootstrap values lower than 100%.

We also found support for a higher abundance of transposable elements (TE) in the W chromosome as compared to all other chromosomes (Fig. 8 & S9). Overall, 45.1% of the cumulative size of the scaffolds assigned to the neoW-W is composed of transposable elements, which represents a 3.31-to 9.95-fold higher content than on the autosomal chromosomes. Interestingly, the TE content over 10-kb windows (Fig. 8) is also much higher on the neoW-4A than on the autosomal 4A chromosome, the other autosomes or, even more interesting, the neoZ-4A region (p<2e-16 for all these comparisons). Even if RepeatModeler was unable to classify most *Z. borbonicus*-specific TE (“no category” in Fig S9), Class I LTR elements seem to have greatly contributed to this higher content in the neoW-W, as well as in the neoW-4A chromosome (Fig S9).

## Discussion

### A new high-quality reference genome for Sylvioidea and *Zosterops*

Using a combination of short Illumina and long PacBio reads, we have generated a high-quality bird assembly of *Z. borbonicus* with a scaffold N50 exceeding one megabase, which is comparable to the best passerine reference genomes available to date (Kapusta & Suh, 2017; Peona et al. 2018). Similar conclusions can be drawn by comparing BUSCO analyses between this assembly and a set of other 26 extant avian genome assemblies (Fig. S1). This assembly is of equivalent quality to the well-assembled congeneric species *Z. lateralis* (Cornetti et al. 2015), thus jointly representing important genomic resources for *Zosterops*, a bird lineage exhibiting one of the fastest rates of species diversification among vertebrates (Moyle et al. 2009).

Avian GC-rich regions are known to be underrepresented in sequencing data because Illumina library construction protocols are biased toward intermediate GC-content (Botero-Castro et al. 2017; Tilak et al. 2018; Peona et al. 2018). Combining moderate coverage (10x) PacBio and Illumina sequencing technologies, we have generated gene models for half of the “hidden” genes of Botero-Castro et al. (2017). The use of long PacBio reads seems therefore a promising solution to partially address the under-representation of GC-rich genes in avian genomes. In the future, it will be interesting to combine long read technologies such as PacBio or Oxford Nanopore with the Illumina library preparation proposed by Tilak et al. (2018).

To further improve the contiguities of the *Z. borbonicus* sequence, we used 26 avian species as pivotal resources to chromosomally organized scaffolds of the *Z. borbonicus* assembly. This strategy has led to a 1.047 Gb chromosome-scale genome sequence for *Z. borbonicus* (Table S1). We were able to obtain assembly statistics comparable to some assemblies using high coverage long-read data (Weissensteiner et al. 2017) or a pedigree linkage map (Kawagami et al. 2014). Importantly, the application of the DeCoSTAR strategy has not only improved the *Z. borbonicus* genome, but also those of 25 other species. Indeed, at the notable exception of the chicken reference genome, all genome assemblies have been improved using DeCoSTAR (4.19-fold improvement of the median scaffold N50), thus demonstrating the utility of the inclusion of other genome assemblies, even fragmented, to polish the genome assembly of a species of interest (Duchemin et al. 2017; Anselmetti et al. 2018).

### Confirming *Zosterops* as great speciators

The availability of the *Z. borbonicus* genome sequence is also an important step to study the evolution of the *Zosterops* lineage, which was described as the “Great Speciator” (Moyle et al. 2009; Cornetti et al. 2015). Indeed, the availability of sequence data for two new species (*Z. borbonicus* and *Z. pallidus*), in addition with the sequence of *Z. lateralis* (Cornetti et al. 2015) helped us to validate the recent origin of this taxon. Moyle et al. (2009) estimated that the white-eyes genus (except *Z. wallacei*) originated in the early pleistocene (∼2.5 Myrs). With more than 80 species, this clade exhibits an exceptional rate of diversification compared to other vertebrates (net rate of diversification without extinction (r): 1-2.5 species per Myr, Magallon and Sanderson, 2001 cited in Harmon, 2018).

The divergence time estimated by Moyle et al. (2009) was based on a biogeographic calibration obtained from the ages of the Solomon Islands. These biogeographic calibrations should however be interpreted with care as previous phylogenetic studies found evidence for older lineages than the emergence ages of the islands for which they are endemic (Heads, 2005; Heads, 2011). This could be the consequence of extinction of mainland relatives leading to long branches of some island species (Heads, 2005; Heads, 2011). More recently, Cai et al. (2019) have obtained similar dates using a larger phylogeny but, again, have applied a biogeographic calibration. In this study, we took the opportunity to combine genetic information of these three *Zosterops* with the other investigated bird species to obtain new estimates using independent fossil calibrations to reassess the conclusions of Moyle et al. (2009) regarding the recent and extensive diversification of *Zosterops*. Using two independent datasets (mitochondrial and nuclear) and methods, we confirmed the recent origin of the genus. Our analyses are consistent with a diversification of white-eyes over less than 5 Myrs and that could be as young as 1 Myr. With the exception of the African Great Lakes cichlids (Genner et al. 2007), the genus *Zosterops* represents one of the most exceptional diversification rates among vertebrates (Lagomarsino et al. 2016). As an example, it is more than ten times higher than the average diversification rate estimated across all bird species by Jetz et al. (2009). Even the large and relatively recent radiation of the Furnariidae (ovenbirds and woodcreepers) has a net rate of diversification markedly lower than the white-eyes (r = 0.16; Derryberry et al. 2010).

### No evidence for a fast-neoZ effect

Taken all together, our analyses are surprisingly consistent with a pattern of substantial reduction of nucleotide diversity, but a low impact of the autosome-to-Z translocation on the molecular evolution of *Z. borbonicus*.

First, using pairwise genome alignments of the zebra finch genome with either the *Z. borbonicus* or the *Z. lateralis* genomes, we found support for a narrow candidate region of 1 kb around position 9.606 Mb of the v.3.2.4 zebra finch 4A chromosome, in which the chromosomal breakpoint likely occurred. This result is consistent and fairly close to the estimate of 10 Mb suggested by Pala and collaborators (2012a) who first demonstrated the translocation of approximately a half of the zebra finch 4A on the Z chromosome using an extended pedigree of the great reed warbler, a Sylvioidea species. A recent article reported a similar estimate (9.6 Mb) in another Sylvioidea species, the common whitethroat (Sigeman et al. 2018). To get this estimate, these latter authors used a very similar approach to ours (H. Sigeman, personal communication).

Second, relative estimates of within-species diversity on both sides of this chromosomal breakpoint (*i.e.* neoZ-4A and neoZ-Z regions) were obtained. As compared to all autosomes, neoZ-4A and neoZ-Z regions of the neo-Z chromosome exhibit reduced levels of within-species diversity in both the ancestral Z chromosome (*i.e.* neoZ-Z:autosomes = 0.605) as well as in the newly translocated region (neoZ-4A:autosomes=0.782), consistent with a substantial loss of diversity associated with this autosome-to-sex transition, following the expected effects of changes in effective population sizes. The neoZ-4A:autosomal-4A nucleotide diversity reported is slightly higher than 0.75 but is in line with a previous report for the common whitethroat (0.82, Pala et al. 2012b). Strongest deviations have however been reported in two other Sylvioidea species, namely the great reed warbler and the skylark (0.15 and 0.42, respectively) but some of the variation might be explained by the moderate number of loci analyzed (Pala et al. 2012b). Sex ratio imbalance or selection are known to contribute to strong deviations from neutral equilibrium expectations of three-fourths (reviewed in Ellegren 2009 and Wilson Sayres, 2018).

Third, we found no support for a fast-Z evolution in the neoZ-4A region, *i.e.* neither an elevated ratio of non-synonymous to synonymous polymorphisms (*π*_*N*_*/π*_*S*_) nor an elevated ratio of non-synonymous to synonymous substitutions (*d /d)* at neoZ-4A genes when compared with autosomal-4A sequences. This result is intriguing as the decrease of nucleotide diversity observed on the neoZ-4A is expected to reflect a decrease in *N*_*e*_ and, therefore, a decrease in the efficacy of natural selection. This should result in an increase of the frequency of slightly deleterious mutations (Ohta 1992; Lanfear et al. 2014). The increase in *d*_*N*_*/d*_*S*_ of avian Z-linked genes compared to autosomes - a pattern that we recovered well in our analyses - has often been interpreted in that way (Mank et al. 2010; Wright et al. 2015). We propose several potential hypotheses to explain the absence of fast-Z on the neoZ-4A regions. First, the hemizygous status of neoZ-4A regions could help to purge the recessive deleterious mutation and, therefore, limit the increase of *π*_*N*_*/π*_*S*_ and *d*_*N*_*/d*_*S*_ as reported in Satyrinae butterflies (Rousselle et al. 2016). Second, the intensity of purifying selection is not only determined by *N*_*e*_ but also by gene expression (Drummond & Wilke 2008; Nguyen et al. 2015) and recombination rate (Hill & Roberston 1966). It is therefore possible that the expression pattern of neoZ-4A genes has changed after the translocation. The change in recombination rate, however, seems to go in the opposite direction as GC and GC* decrease in the neoZ-4A region, suggesting a decrease in recombination rate and, therefore, a decrease in the efficacy of natural selection.

Finally, base composition has changed after the translocation to the Z chromosome. We indeed found evidence for lower GC content and GC* in the neoZ-4A region than in the remaining autosomal region of the 4A chromosome. In birds, as in many other organisms, chromosome size and recombination rate are negatively correlated (Backström et al. 2010), probably because one recombination event occurs per chromosome arm per meiosis. Given that GC content strongly negatively correlates with chromosome size (Eyre-Walker, 1993; Pessia et al. 2012), the observed difference in GC content is a likely consequence of changes in the intensity of the recombination-associated effect of gBGC (Duret & Arndt, 2008; Duret & Galtier, 2009), resulting from the instantaneous changes in chromosome sizes due to the translocation. From the translocated region of the ancestral 4A chromosome point of view, the chromosomal context has drastically changed from an ancestral ∼21 Mb 4A chromosome to a ∼90 Mb neo-Z chromosome probably with a lower recombination rate. Similarly, from the remaining 4A chromosome point of view, the chromosomal context has drastically changed too, from a ∼21-Mb to a ∼11-Mb chromosome. As a consequence, we can expect that base composition has evolved in the opposite direction with an increase in GC. However, this hypothesis must be qualified since we were unable to find any change in GC* at autosomal 4A genes, suggesting that the chromosome-scale recombination rate might not have been impacted by the translocation. This could be possible if the translocation occurred close to the centromere (*i.e.* a nearly whole-arm translocation). In this case, the overall recombination rate is not expected to change.

### Convergent chromosomal footprints of a neoW degeneration

Degeneration of the non-recombining chromosome, *i.e.* the chromosome only present in the heterogametic sex, has been explored in depth in a variety of species (*e.g.* Bachtrog & Charlesworth, 2002; Papadopulos et al. 2015). Long-term gradual degeneration through the accumulation of deleterious mutations, partial loss of adaptive potential and gene losses are expected to start soon after species cease to recombine due to a series of factors including Muller’s ratchet, linked selection and the Hill-Robertson effect (Charlesworth & Charlesworth, 2000; Bachtrog, 2005; Sun & Heitman 2012).

To investigate this degeneration, we have first identified 218 scaffolds with a female-specific pattern in read coverage, for a total of 7.1 Mb. Among these scaffolds, we have assigned 174 scaffolds to the neoW-4A region, because of a high level of homology with the zebra finch 4A chromosome, thus representing a neoW-4A of a total length of 5.48 Mb. The absence of any neoW scaffold homologous to the 4A *T. guttata* chromosome between positions 1,756,080 and 3,509,582 suggests a large chromosomal deletion. The total length of the neoW-W region we have assigned is low, only representing 1.23 Mb of sequence, which is far from the 6.94 Mb sequence that Smeds and collaborators (2015) identified in the collared flycatcher genome or the 6.81 Mb sequence reported in the reference Chicken genome (GRCg6a version, International Chicken Genome Sequencing Consortium 2004; Warren et al. 2017). It is however important to specify that our objective was not to be exhaustive, but rather to focus on the longest scaffolds for which both estimates of the median coverage and alignments against the zebra finch chromosome 4A were considered reliable enough to be confident in their assignment to the W chromosome, particularly in a context of the intense activity of transposable elements. Given the known difficulty to sequence and assemble the W chromosome (*e.g.* Tomaszkiewicz et al. 2017), such a reduced-representation of the W chromosome was expected. Our intent was to get sufficient information to study the molecular evolution of neoW-W specific genes too. Obtaining a high-quality sequence of the neoW chromosome for the *Z. borbonicus* species, while possible, would require a considerable additional sequencing effort to be achieved.

Then, by aligning *Z. borbonicus* assembly against the zebra finch 4A chromosome, we found support for a candidate scaffold supporting the chromosomal breakpoint. Alignments on both ends of this scaffold suggest a potential chromosomal breakpoint occurring around positions 9.603-9.605 Mb of the v.3.2.4 zebra finch 4A chromosome, which is remarkably close to our estimate for the translocation on the neoZ-4A. Importantly, such an observation therefore supports an evolution of neoW-4A and neoZ-4A regions from initially identical gene sets. Interestingly, we have also identified a large chromosomal deletion on the W chromosome, which represents another expected early signature of the W degeneration (Charlesworth, 1991).

We have found support for a highly reduced level of nucleotide diversity in the neoW chromosome as compared to autosomes. This also holds true for the neoW-4A region (mean neoW-4A:autosomal nucleotide diversity = 0.36), which is in broad agreement with the hypothesis of a three-fourth reduction in effective population size associated to an autosomal-to-W or autosomal-to-Y translocation. Our overall result of low within-species diversity on the W chromosome is however not as drastic as compared with the dramatically reduced diversity observed by Smeds and collaborators (2015) on the W chromosome of several flycatcher species, with a W:autosomal diversity ranging from 0.96% to 2.16% depending on populations and species. Non-recombinant W chromosome of *Z. borbonicus* also exhibits elevated π_N_/π_S_ in the neoW-W (0.78) as well as in the neoW-4A regions (0.42), representing 3.68-fold and 1.97-fold higher ratios than for the autosomal genes, respectively (4.59-fold and 2.46-fold higher ratios when compared with autosomal-4A genes only). Higher d_N_/d_S_ at neoW-4A genes as compared to their neoZ-4A gametologs were also observed. This higher d_N_/d_S_ ratio is in agreement with the pattern observed in another Sylvioidea species, the common whitethroat (Sigeman et al. 2018) and, more broadly, with other young sex chromosome systems (*e.g.* Marais et al. 2008). Sigeman et al. (2018) also reported an association between amino acid and gene expression divergences for the neoW-4A. Altogether, these results are consistent with an accumulation of deleterious mutations associated with the strong reduction of the net efficacy of natural selection to purge deleterious mutations (Charlesworth & Charlesworth, 1997).

TE accumulation on W or Y chromosome is suspected to play a particularly important role in the first phases of the evolution of chromosome differentiation (Bachtrog, 2003). To investigate this, we have *de novo* identified *Z. borbonicus*-specific TE and have analyzed distribution and abundance of TEs. This has led to the identification of a high TE load in the ancestral W chromosome (45.1%), which is the same order of magnitude as the reported value for the W chromosome of *Ficedula albicollis* (48.5%, Smeds et al. 2015) or *Zonotrichia albicollis* (51.1%, Davis et al. 2010). Interestingly, we found support for a high TE load in the translocated region too (21.6%), which is approximately twice the observed TE content of any other autosomal chromosome, including the autosomal 4A chromosome. TE classification, albeit incomplete, supports an important contribution of class I LTR elements to the overall TE load. LTR elements seem to be particularly active in the zebra finch (Kapusta and Suh, 2017) or in the collared flycatcher genomes (Suh et al. 2018) suggesting that the recent burst of LTR elements on autosomes may have facilitated the accumulation of TEs on the neoW-4A chromosomes. The most probable hypothesis is that LTR elements are particularly retained in low-recombination regions. Following this hypothesis, the non-recombining neoW-4A region might therefore be viewed as an extreme case in terms of the retention of these TE insertions. Although much more data and work will be needed in the future to analyze in greater depth this accumulation of TEs, and particularly the determinants of this accumulation, our results suggest that LTR elements may have virtually played a major role in altering gene content, expression and/or chromosome organization of the newly translocated region of the Sylvioidea neo-W chromosome.

## Conclusions

In this study, we generated a high quality reference genome for *Z. borbonicus* that has provided us unique opportunities to investigate the molecular evolution of neo-sex chromosomes, and more broadly to improve our understanding of avian sex chromosome evolution. Since this species belong to the Sylvioidea, one of the three major clades of passerines, comprising close to 1,300 species, we can reasonably anticipate that this chromosomal-scale assembly will serve as a reference for a large diversity of genome-wide analyses in the Sylvioidea lineage itself, and more generally in passerine birds. Interestingly, Sylvioidea is becoming an important animal taxon for the study of sex chromosomes (Dierickx et al. 2019; Sigeman et al. 2019).

Through detailed analyses of the evolution of the newly sex chromosome-associated regions, we found evolutionary patterns that were largely consistent with the classic expectations for the evolution of translocated regions on sex chromosomes, including evidence for reduction of diversity, ongoing neoW chromosome degeneration and base composition changes. A notable exception was the neoZ region for which no fast-neoZ effect was identified. Although most of the analyses are congruent, our report is based on a limited number of individuals and from only one population of *Z. borbonicus*. Further investigations based on a complementary and extensive dataset will probably help to fine-tune the conclusions, especially regarding the lack of any fast-Z effect or the drastic increase of TE on the neoW-4A. Lastly, given the huge difference in diversity between autosomes and sex chromosomes, we emphasize the importance of taking into account sex chromosomes for local adaptation studies, at least by scanningn autosomes and sex chromosomes separately (see Bourgeois et al. 2018 for an example).

## Materials & Methods

### DNA and RNA extraction, sequencing

We extracted DNA from fresh tissues collected on a *Z. borbonicus* female individual (field code: 15-179), which died accidentally during fieldwork in May 2015 at Pas de Bellecombe (Gîte du volcan, Réunion; coordinates: S: −21.2174, E: 55.6872; elevation: 2246m above sea level). Sampling was conducted under a research permit (#602) issued to Christophe Thébaud by the Centre de Recherches sur la Biologie des Populations d’Oiseaux (CRBPO) – Muséum National d’Histoire Naturelle (Paris).

We also extracted DNA of a *Zosterops pallidus* female collected in February 2015 in South Africa, Free state province, Sandymount Park, 10 kms from Fauresmith (coordinates: S: - 29.75508, E: 25.17733). The voucher is stored at the Museum National d’Histoire Naturelle (MNHN), Paris, France, under the code MNHN ZO 2015-572 and a tissue duplicate is deposited in the National Museum Bloemfontein (South Africa).

For both samples, 9µg of total genomic DNA were extracted from liver and/or muscle, using DNeasy Blood and Tissue kit (QIAGEN) following the manufacturer instructions. Each of these samples was sequenced as followed: one paired-end library with insert sizes of 300bp and three mate-pair libraries (3 kb, 5 kb and 8 kb) using Nextera kit. Libraries and sequencing were performed by platform “INRA plateformes Génomes et transcriptomes (GeT-PlaGe)”, Toulouse, France. Illumina sequencing was also performed at the platform GeT-PlaGe using Illumina HiSeq 3000 technology.

To improve genome assembly of Z. *borbonicus*, an additional sequencing effort was made by generating 11X coverage of PacBio long reads data. 20µg of high molecular weight DNA were extracted from muscle using MagAttract HMW DNA kit (QIAGEN) following manufacturer instructions. The PacBio librairies and sequencing were performed at Genome Québec (Centre d’innovation Génome Québec et Université McGill, Montréal, Quebec, Canada) using a PacBio RS platform.

After the accidental death of the *Z. borbonicus*, the brain of the freshly dead bird was extracted and then stabilized using RNAlater (Sigma). Total RNA of *Z. borbonicus* individual was extracted from the dissected tissue sampleusing RNeasy Plus Mini Kit (Qiagen) following manufacturer’s instructions (RNA integrity number: 7.9). Both the RNAseq library preparation and the Hiseq2500 sequencing (1 lane) were performed at the genomic platform MGX-Montpellier GenomiX.

### Genome assembly

The paired-end reads were filtered using Trimmomatic (v0-33; Bolger et al., 2014) using the following parameters: “ILLUMINACLIP:TruSeq3-PE-2.fa:2:30:10 LEADING:3 TRAILING:3 SLIDINGWINDOW:4:15 MINLEN:50”. The mate-pair reads were cleaned using NextClip (v1.3.1; Leggett et al. 2013) using the following parameters: “--min_length 20 -- trim_ends 0 --remove_duplicates”.

Paired-end and mate-pair reads were assembled using SOAPdenovo (v2.04; Luo et al 2012) with parameters “-d 1 -D 2”. Several k-mers (from 27 to 37-mers) were tested and we chose the assembly maximizing the N50 scaffold length criteria. Next, we applied Gapcloser v1.10 (a companion program of SOAPdenovo) to fill the gap in the assemblies.

Given that the PacBio technology produces long reads but with quite high sequencing error rates, we used LoRDEC (v0.6; Salmela & Rivals 2014) to correct the PacBio reads of the *Z. borbonicus* individual, using the following parameters: “-k 19 -s 3”. In brief, LoRDEC corrects PacBio reads (both insert/deletions and base call errors) by the use of Illumina paired-end reads, a technology producing short reads only, but with a much higher base call accuracy and depth of coverage. The corrected PacBio reads were then used to scaffolds the SOAPdenovo assembly using SSPACE-LongRead (v1.1; Boetzer & Pirovano 2014). For *Z. borbonicus*, we also used MaSuRCA (v3.2.4; Zimin et al. 2017) to perform a hybrid assembly with a mixture of short and long reads. MaSuRCA produced an assembly with similar quality but slightly shorter than SOAPdenovo+SSPACE-LongRead (Table S1).

Several statistics were computed using assemblathon_stats.pl script (Bradnam et al. 2013; https://github.com/ucdavis-bioinformatics/assemblathon2-analysis) to evaluate the different genome assemblies. These statistics include genome size, number of scaffolds, scaffold N50, scaffold N90 and the proportion of missing data (N%). Additionally, we used BUSCO (v3.0.2; Waterhouse et al. 2017), a commonly used tool for evaluating the genome completeness based on the content in highly conserved orthologous genes (options: “-m genome -e 0.001 -l aves_odb9 -sp chicken”).

The mitochondrial genome was assembled using mitobim (v1.8; Hahn et al. 2013) with the mitochondrial genome of *Z. lateralis* (accession: KC545407) used as reference (so-called “bait”). The mitochondrial genome was automatically annotated using the web server mitos2 (http://mitos2.bioinf.uni-leipzig.de; Bernt et al. 2013). This annotation was manually inspected and corrected using alignment with the other *Zosterops* mitochondrial genomes available in genbank.

### Genome-guided De novo Transcriptome Assembly

RNA-seq reads were used to generate a transcript catalogue to train the gene prediction software. First, RNA-seq reads were filtered with trimmomatic (v0-33; Bolger et al., 2014) using the following parameters: “ILLUMINACLIP:TruSeq3-PE.fa:2:30:10 SLIDINGWINDOW:4:5 LEADING:5 TRAILING:5 MINLEN:25”. Second, the filtered RNA-seq reads were mapped onto the reference genome using HISAT2 (Kim et al. 2015). HISAT2 performed a splice alignment of RNA-Seq reads, outperforming the spliced aligner algorithm implemented in TopHat (Kim et al. 2015). Finally, two methods were used to assemble the transcripts from the HISAT2 output bam file: i) Cufflinks (Trapnell et al. 2010) using the following parameters: “-q -p 10 -m 300 -s 100” and ii) Trinity (v2.5.0; Haas et al. 2013) using the following parameters:

“--genome_guided_max_intron 100000”.

### Protein-coding Gene Annotation

Gene annotation was performed using the PASA pipeline combined with EVidenceModeler (Haas et al. 2008; Haas et al. 2011; https://github.com/PASApipeline/PASApipeline/wiki). The complete annotation pipeline involved the following steps:

1. *ab initio* gene finding with Augustus (Stanke and Waack 2003; http://bioinf.uni-greifswald.de/augustus/) using the parameter: “- -species=chicken”.
2. Protein homology detection and intron resolution using genBlastG (She et al. 2011; http://genome.sfu.ca/genblast/download.html). Protein sequences of several passerines were used as references, namely zebra finch (*Taeniopygia guttata*; assembly taeGut3.2.4; Warren et al. 2010), collared flycatcher (*Ficedula albicollis*; assembly FicAlb_1.4; Ellegren et al. 2012), medium ground-finch (*Geospiza fortis*; assembly GeoFor_1.0; (Zhang et al. 2012) and hooded crow (*Corvus cornix*; accession number JPSR00000000.1; Poelstra et al. 2014). Genblast parameters were: “-p genblastg -c 0.8 -r 3.0 -gff - pro -cdna -e 1e-10”.
3. PASA alignment assemblies based on overlapping transcript from Trinity genome-guided *de novo* transcriptome assembly (see above) (Haas et al. 2003). This step involved the so-called PASAPipeline (v2.2.0; https://github.com/PASApipeline/PASApipeline/) used with the following parameters: “-C -r -R -- ALIGNERS blat,gmap”.
4. Next, EVidenceModeler (v1.1.1; Haas et al. 2008) was used to compute weighted consensus gene structure annotations based on the previous steps (1 to 3). We used the following parameters: “--segmentSize 500000 --overlapSize 200000” and an arbitrary weight file following the guidelines provided at http://evidencemodeler.github.io/.
5. Finally, we used the script “pasa_gff3_validator.pl” of PASA add UTR annotations and models for alternatively spliced isoforms.

Finally, StringTie (Pertea et al. 2015) was also used to estimate the proportion of annotated CDS with RNA-seq information support.

We defined three sets of genes depending on their reliability, namely the “high reliability” set corresponding to genes with a low TE content in coding regions (<10%, Fig. S2) and with at least a transcript support (“high”), the “moderate reliability” set corresponding to genes with either a low TE content (<10%) or with at least a transcript support (“moderate”) and a “low reliability” set containing the remaining genes.

### Orthology detection

We used the available passerine genomes (namely, zebra finch, collared flycatcher, white-throated sparrow and hooded crow) plus the high-coverage genomes (>100x) of Zhang et al. (2014) (Table S2). Orthology detection was performed using OrthoFinder (v2.2.6; Emms & Kelly 2015). Single copy (one-to-one) orthologs were extracted from OrthoFinder results to perform multi-species alignment with TranslatorX (v1.1; Abascal et al. 2010) using MAFFT (Katoh et al. 2002) to build the alignment. Alignments were inspected by HMMCleaner (v1.8; Amemiya et al. 2013; Philippe et al. 2017) to exclude badly aligned sites. Next, dubious, highly divergent, sequences were excluded using trimAl (Capella-Gutiérrez et al. 2009) option “-resoverlap 0.60 -seqoverlap 80”. We also used genome assemblies of three passerine species for which no gene annotation sets were publicly available, namely the silvereye, *Z. lateralis* (Cornetti et al. 2015), the Orange River white-eye *Z. pallidus* (this study) and the willow warbler *Phylloscopus trochilus* (Lundberg et al. 2017). For these genomes, gene orthology detection was conducted using AGILE (Hughes & Teeling 2018). AGILE is a pipeline for gene mining in fragmented assembly overcoming the difficulty that genes could be located in several scaffolds. We applied AGILE using *Z. borbonicus* single copy orthologs as query genes.

### Filtering scaffolds originating from autosomes, W and Z chromosomes

Given that we have sequenced a female genome, we likely assembled contigs from W, Z and autosomal chromosomes. To identify scaffolds originating from sex chromosomes or autosomes, we mapped seven females and five males reads from *Z. borbonicus* birds published in Bourgeois et al. (2017) onto our female genome assembly. We first mapped all raw reads against the *Z. borbonicus* reference genome using BWA mem v. 0.7.5a (Li, 2013), we then removed duplicates with Picard 1.112 (http://broadinstitute.github.io/picard). We then used Mosdepth (Pedersen & Quinlan, 2018) to compute median per-site coverage for each scaffold, following the same strategy than in Smeds et al. (2015). For each *i* scaffold, total coverage for males and females were computed as the sum of coverage of all individuals. Then, we compute a normalized coverage per scaffolds for male as:

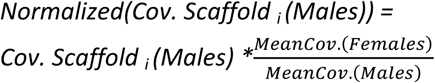

Where “*Mean Cov. (Females)*” and “*Mean Cov. (Males)*” corresponds to the median coverages for males and females across all scaffolds longer than 1 Mb. This normalization is intended to take into account the different number of male and female individuals.

Next, we used the normalized median per-site coverage to detect W-linked scaffolds (zero and above 5X in males and females, respectively), Z-linked (female with less than 0.75 the male coverage and male with > 9X coverage) and autosomes (all the remaining scaffolds corresponding female and male with a roughly similar median coverage). To decrease the probability of false identification due to the mapping in repeat-rich regions, this approach has only been performed using coverage data from scaffolds longer than 10 kb (3,443 / 97,503 scaffolds, >96% of the assembly size).

### Pseudo-chromosome assembly

We first aligned soft-masked *Z. borbonicus* scaffolds on the soft-masked Zebra Finch genome (*Taeniopygia guttata*) using LASTZ v. 1.04.00 (Schwartz et al. 2003). For *Z. borbonicus, de novo* transposable elements prediction were performed using Repeat Modeler v. 1.0.11 and genome assembly soft-masking using Repeat Masker v. 4.0.3 (Smit et al. 2013-2015). For *T. guttata*, we used the soft-masked genome v. 3.2.4 (taeGut3.2.4) made available by the Zebra Finch genome consortium. This soft-masking procedure was put in place in order to exclude these regions for the LASTZ’s seeding stage, and thereby to avoid finding alignments in these highly repeated regions. We then filtered LASTZ alignments hits in order to only keep reliable hits (5% longest hits and with sequence identities greater than the median over all alignments). For each scaffold, we then defined syntenic blocks as adjacencies of several reliable hits. A syntenic block represents a homologous region between zebra finch and *Z. borbonicus* starting at the first reliable hit and ending at the last one. Syntenic blocks covering at least 80% of a *Z. borbonicus* scaffold size and 80-120% the corresponding homologous region in the Zebra Finch genome were automatically anchored to its *T. guttata* chromosome position, assuming complete synteny between the zebra finch and *Z. borbonicus*. Unanchored scaffolds were then manually identified by visually inspecting the summary statistics and positions of all raw alignments. In case of chimeric scaffolds, these scaffolds were cut into two or several new scaffolds assuming that this chimerism is due to a contigging or a scaffolding artifact.

Second, we used DeCoSTAR, a computational method tracking gene order evolution from phylogenetic signal by inferring gene adjacencies evolutionary histories, in order to not only improve the genome assembly of *Z. borbonicus*, but also 26 additional extant avian assemblies including the high coverage (>100X) of Zhang et al. 2014 and the other passerine genomes (Table S2). The gene trees were obtained for phylogenies of single-copy orthologs (see above) estimated using IQ-TREE model GTR+G4 and 1000 so-called “ultra-fast bootstrap” (v1.6.2; Nguyen et al. 2014). A snakemake pipeline (https://github.com/YoannAnselmetti/DeCoSTAR_pipeline) were used to apply DeCoSTAR on gene orders and phylogenies.

Finally, scaffold orders given by LASTZ and DeCoSTAR were manually reconciled to determine the consensus scaffold orders along the *Z.borbonicus* chromosomes.

### Molecular dating

We performed two independent molecular dating analyses. First, we used relaxed molecular clock analysis based on nuclear sequences and fossil calibrations applied on the full bird phylogeny. Second, we applied the approach of Nabholz et al. (2016) using white-eyes body mass to estimate their mitochondrial substitution rate.

In the first approach, molecular dating analyses were performed using the 27 species selected for the DeCoSTAR analysis plus the silvereye, the Orange River white-eye and the willow warbler leading to a total of 30 species. We restricted our analyses to single-copy orthologs with low GC content. These genes are known to evolve slowing and more clock-like than GC rich genes (Nabholz et al. 2011; Jarvis et al. 2014). To do that, we randomly selected 100 single-copy orthologs coded by chromosome 1 and 2 excluding genes at the beginning and at the end of the chromosomes (minimum distance from each chromosome tip based on the *T. guttata* genome: 10 Mbp). This step was replicated ten times to evaluate the robustness of the inferences among different sets of genes.

We used several fossil calibrations sets summarized in Table S3. Our different calibration sets reflect the current uncertainty surrounding bird diversification dates. We used a conservative set with only one maximum bound at 140 Myrs for the origin of Neornithes (set 4). At the other end, we used another set with a narrow constraint between 28 and 34 Myrs for the Suboscines / Oscines divergence (set 3). This constraint could turn out to be incorrect with future advances in the bird fossil records. For all the calibration sets, the Neognathae / Palaeognathae divergence minimal age was set to 66 Myrs using *Vegavis iaai* fossil (Benton et al. 2009; Mayr, 2013; Ksepka & Clarke, 2015). The maximum age of this node is much more difficult to select. We have opted for two maximal ages. The first one at 86.5 Myrs following the rationale of Prum et al. (2015) based of the upper bound age estimate of the Niobrara Formation (set 1, 2 and 3). We used another very conservative maximum bound at 140 Myrs (set 4). This is the maximum age estimated for the origin of Neornithes by Lee et al. (2014) using an extensive morphological clock analysis. Additionally, in sets 1 and 3, we constrained the divergence between Passeriformes / Psittaciformes to be between 53.5 and 65.5 Myrs, assuming the complete absence of passerine bird species during the Cretaceous. In sets 2 and 4, we only used a minimum bound on the first fossil occurrence of passerines in the Eocene (Boles, 1997) and on the stem Psittaciformes fossil *Pulchrapollia gracilis* (Dyke & Cooper, 2000). In calibration set 3, we constrained the Oscines/Suboscines split between 28 and 34 Myrs, following the rationale of Mayr (2013) assuming crown Oscines and Suboscines originated in the early Oligocene (28 Myrs, Mayr & Manegold 2006) and based on the absence of Eocene fossil records discovered so far (Eocene/Oligocene limit is at 34 Myrs). All the other minimum-bound calibrations we used followed the suggestions of Ksepka & Clarke (2015) and are presented in Table S3.

Molecular dating was performed using Phylobayes (v4.1; Lartillot et al. 2009) using a CAT-GTR substitution model. For the relaxed clock model, we used the log-nomal auto-correlated rates (ln) model. We also tested the uncorrelated gamma multipliers (ugam) model that gave similar results than the ln model (results not shown). We used uniform prior on divergence times, combined with soft calibrations (Rannala and Yang, 2007; Yang and Rannala, 2006). The MCMC were run for at least 8,000 cycles. MCMC convergence were diagnosed by running two independents MCMC and by visually checking the evolution of the likelihood and other parameters along the Markov chain (in “.trace” files).

Additionally, we applied an independent molecular dating using the method proposed by Nabholz et al. (2016). Seven complete mitochondrial *Zosterops* genomes were downloaded from Genbank, including *Z. erythropleuros* (KT194322), Z. japonicus (KT601061), *Z. lateralis* (KC545407), *Z. poliogastrus* (KX181886), *Z. senegalensis* (KX181887), *Z. senegalensis* (KX181888). We also included the sequences of the Réunion grey white-eye and the Orange River white-eye assembled in the present study. The mitochondrial sequences of *Yuhina diademata* (KT783535) and *Zoothera dauma* (KT340629) were used as outgroups. Body mass for all *Zosterops* species were obtained from Dunning (2007) and the median of these body masses was computed. Phylogenetic relationship and branch length were estimated using IQ-TREE with a HKY+G4 substitution model using third codon positions only. Then, we applied the formula of Nabholz et al. 2016 to derive the substitution rate (substitution per site per Myr) as 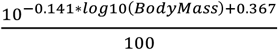 and 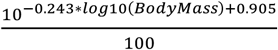 for the minimal and maximal rate where “*Body Mass*” is the median body mass in grams (logarithm of base 10). Next, we computed the median divergence time between the silver-eye (*Z. lateralis)* and all the other white-eye species to obtain an estimate of the crown *Zosterops* clade age. Finally, we divided this divergence time by the rate obtained with the formula above to obtain divergence dates in Myrs.

### Genome-wide estimates of nucleotide diversity

Based on the previously generated BAM files (see ‘*Filtering scaffolds originating from autosomes, W and Z chromosomes*’ section), we then use GATK to generate the gvcf from the six parents of the three progenies described in Bourgeois et al. 2017 (three males and three females). Joint genotyping, as well as all subsequent analyses, was then performed separately for the three males and the three females. We followed all the GATK best practices under the GATK suite, except for variant filtration which was performed using a custom script to speed up computations. This step however followed the same procedures than under GATK, assuming the following thresholds: QD>2.0, FS<60.0, MQ>40.0, MQRANKSUM>-2.0, READPOSRANKSUM>-2.0 and RAW_MQ > 45000.

For each individual, we then reconstructed two genomic sequences. At each position of the genome, the position (reference or alternate if any) was added to the sequence if the coverage at the position was between 3 and 50. In any other case, a “N” character was added in order to keep the sequence length the same. We then computed the Tajima’s π estimator of nucleotides diversity and the GC content over non-overlapping 10 kb windows. To be highly conservative in the analysis of the neoW chromosome, all windows associated to W scaffolds and found covered in the male (ZZ) dataset were excluded from the analysis of the female (ZW) dataset, even if this non-zero coverage was observed at a single base over the genomic window. Such a non-zero coverage is expected caused by read mismapping, particularly in TE regions.

### Estimates of non-synonymous and synonymous divergences and GC equilibrium

For the neoZ-4A and neoW-4A, we made a specific effort to retrieve the paralogous sequences. Assuming that the two translocated regions evolved from the same gene sets (see results), each gene located on the neoZ-4A are expected to have a paralog on the neoW-4A (so called “gametologs”, Pala et al. 2012a,b). As a consequence, most gametologs have been eliminated during our selection of the single-copy orthologs. We visually inspected all the alignments containing a neoZ-4A gene and then tried to identify the corresponding copy in a scaffold assigned to the neoW chromosome.

Given the drastically different number of genes between autosomes, neoZ-Z and neoZ-4A regions, we subsampled the data to match the category with the lowest number of genes (i.e., the neoZ-4A region). Then, we concatenated genes and computed *d*_*N*_ and *d*_*S*_ as the sum of non-synonymous and synonymous branch lengths respectively. This will decrease the variance of the estimated *d*_*N*_*/d*_*S*_ ratios and limit problems associated to low *d*_*S*_ values (Wolf et al. 2009). Finally, the variability in *d*_*N*_*/d*_*S*_ was evaluated by bootstrapping genes 1000 times within each genomic region. For GC equilibrium (GC*), we used the nonhomogeneous model T92 (Galtier and Gouy 1998) implemented in the BPPSUITE package (Dutheil and Boussau 2008, http://biopp.univ-montp2.fr/wiki/index.php/BppSuite) based on the Bio++ library (Guéguen et al. 2013) to infer GC* at third codon position for each branch. We used a similar bootstrapping strategy (1000 times within each genomic region) to evaluate the variation in GC*.

For the comparison between neoZ-4A and neoW-4A, we used the *d*_*N*_*/d*_*S*_ and GC* of neoZ-4A and neoW-4A genes for the branch leading to Z. borbonicus. When the neoW-4A sequences of Z. borbonicus was very closely related to a sequence of Z. pallidus, we assumed that the Z. pallidus sequence also came from the neoW-4A regions and we computed the *d*_*N*_*/d*_*S*_ of the ancestral branch of these two species. We evaluated the different in *d*_*N*_*/d*_*S*_ using a paired-samples Wilcoxon test (also known as Wilcoxon signed-rank test). The *d*_*N*_*/d*_*S*_ was estimated using CODEML (Yang 2007) using a free-ratios model (model = 2). We also checked for the presence of frameshift and premature stop codon within the neoW-W sequenced using macse (v2, Ranwez et al. 2007).

### GC and TE contents

We used the automated approach implemented in RepeatModeler Open (v.1.0.11, Smit & Hubley, 2008) to *de novo* detect *Z.borbonicus*-specific TE consensus. The generated list of *de novo* TE sequences was merged to the chicken repetitive sequences publicly available in Repbase (Jurka et al. 2005). We then used this set of sequences as a custom library for RepeatMasker (v.open-4.0.3, Smit et al. 2013) to generate a softmasked version of the *Z. borbonicus* genome assembly. Then, we used a non-overlapping 10-kbp sliding windows approach to calculate the GC and TE contents along the whole genome. All genomic windows composed of more than 50% of Ns were excluded to ensure accurate estimates of local TE and GC contents.

All statistical analyses were performed using R v. 3.4.4 (R core Team, 2018). Some analyses and graphics were performed using several additional R packages: APE (Paradis & Strimmer, 2004), beanplot (Kampstra, 2008), circlize (Gu et al. 2014), cowplot (Wilke, 2016), ggplot2 (Wickham, 2016), phytools (Revell, 2012) and plotrix (Lemon, 2006).

## Data availability

Nuclear and mitochondrial genome sequences, scripts and programs used are available at the following FigShare repository: https://figshare.com/s/122efbec2e3632188674. Raw reads and genome sequences are available on SRA and GenBank (Bioproject: PRJNA530916). Mitochondrial genome sequences are available on GenBank (accession numbers: MK524996, MK529728).

## Acknowledgments

This research was funded by the French ANR (BirdIslandGenomic project, ANR-14-CE02-0002). The analyses benefited from the Montpellier Bioinformatics Biodiversity (MBB) platform services and the genotoul bioinformatics platform Toulouse Midi-Pyrenees (Bioinfo Genotoul). We are grateful to Jérôme Fuchs for providing the mitochondrial alignments, to Fabien Condamine for constructive discussions on molecular dating and Anna-Sophie Fiston-Lavier for providing advice on the analysis of transposable elements. We thank the Reunion National Park for granting us permission to conduct fieldwork in Pas de Bellecombe, Reunion. We are grateful for the logistic support provided by the field station of Marelongue, funded by the P.O.E., Reunion National Park and OSU Reunion. We are grateful to the provincial authorities in the Free State (South Africa) for granting permission to collect samples and specimens (permit 01-24158) and to Dawie de Swardt (National Museum Bloemfontein) for help with organizing field work.

This preprint has been peer-reviewed and recommended by Peer Community In Evolutionary Biology (https://doi.org/10.24072/pci.evolbiol.100073) We are grateful to the PCI recommender Kateryna Makova as well as three reviewers (Melissa Wilson, Gabriel Marais and an anonymous reviewer) for providing excellent reviews based on a previous version of the manuscript.

## Supplementary Information

**Table S1:**
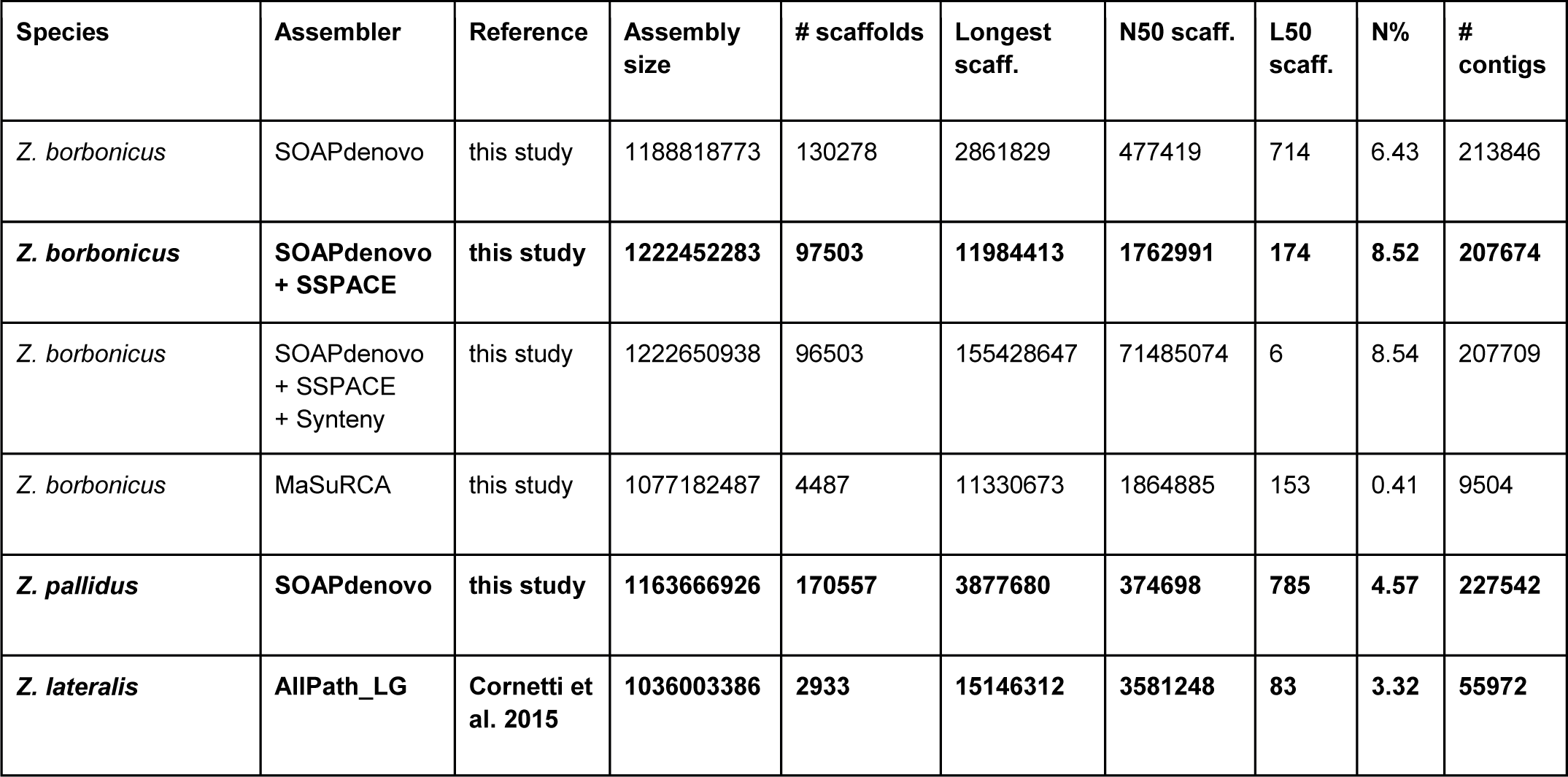
Summary statistics of the *Zosterops* genome assemblies as computed by Assemblathon2. Values in bold correspond to the summary statistics of the publicly available sequences.

| Species | Assembler | Reference | Assembly size | # scaffolds | Longest scaff. | N50 scaff. | L50 scaff. | N% | # contigs |
| --- | --- | --- | --- | --- | --- | --- | --- | --- | --- |
| *Z. borbonicus* | SOAPdenovo | this study | 1188818773 | 130278 | 2861829 | 477419 | 714 | 6.43 | 213846 |
| ***Z. borbonicus*** | **SOAPdenovo + SSPACE** | **this study** | **1222452283** | **97503** | **11984413** | **1762991** | **174** | **8.52** | **207674** |
| *Z. borbonicus* | SOAPdenovo + SSPACE + Synteny | this study | 1222650938 | 96503 | 155428647 | 71485074 | 6 | 8.54 | 207709 |
| *Z. borbonicus* | MaSuRCA | this study | 1077182487 | 4487 | 11330673 | 1864885 | 153 | 0.41 | 9504 |
| ***Z. pallidus*** | **SOAPdenovo** | **this study** | **1163666926** | **170557** | **3877680** | **374698** | **785** | **4.57** | **227542** |
| ***Z. lateralis*** | **AIPath_LG** | **Cornetti et al. 2015** | **1036003386** | **2933** | **15146312** | **3581248** | **83** | **3.32** | **55972** |

**Table S2:**
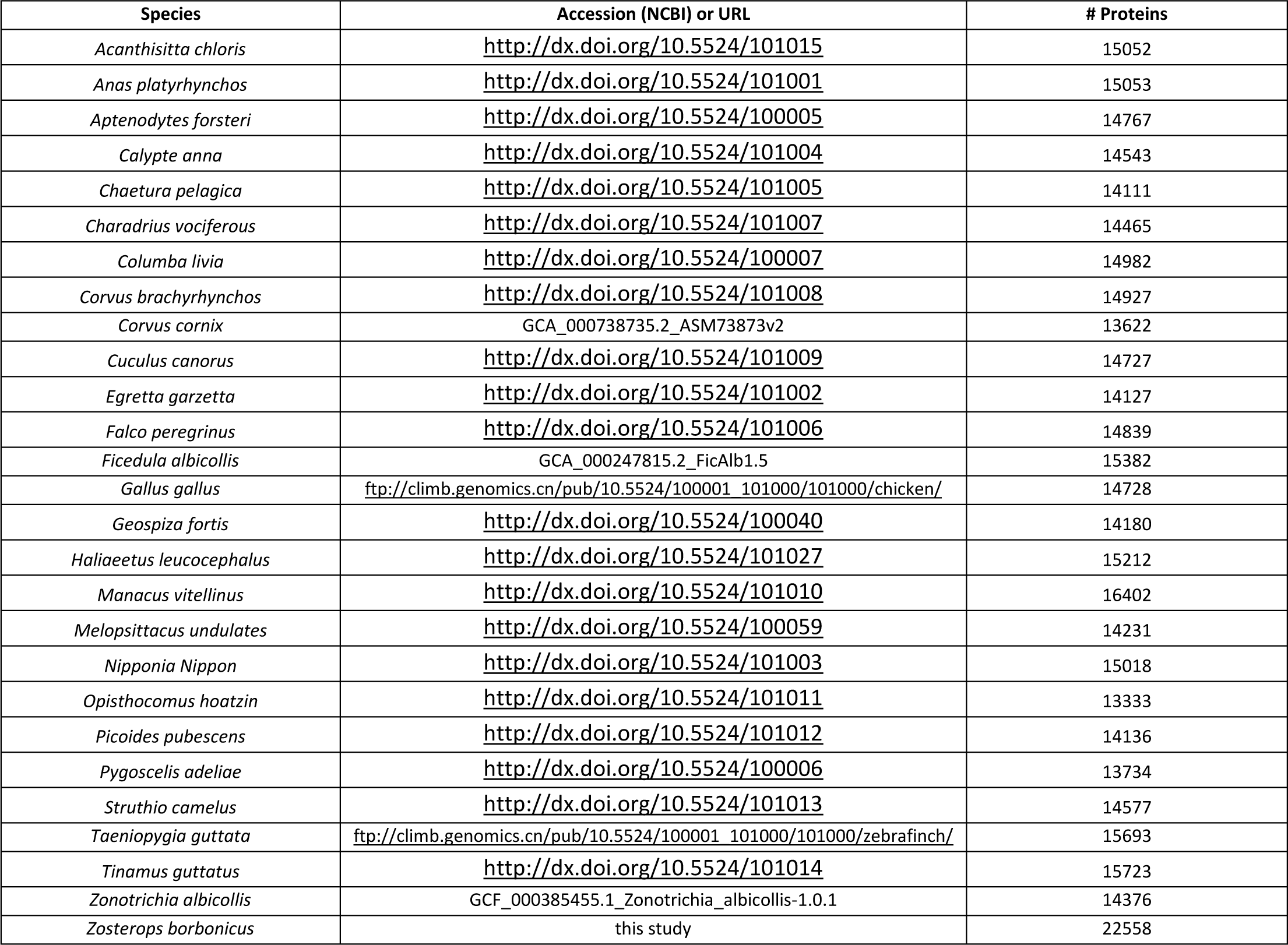
List of the 27 avian species used in DeCoSTAR. Number of CDS translated to protein and used for orthology detection.

**Table S3:**
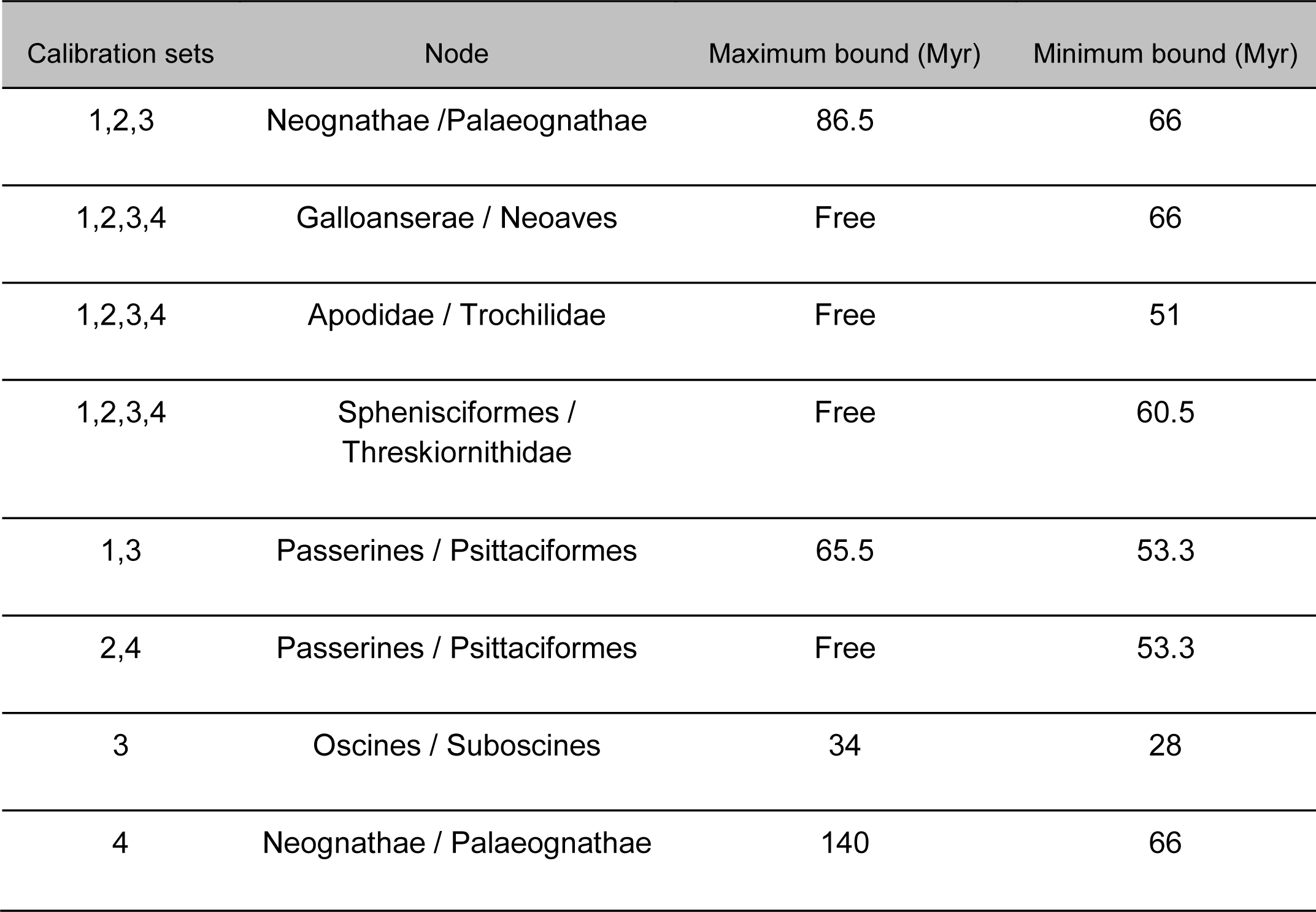
Fossil calibration combinations used in the molecular dating analyses

| Calibration sets | Node | Maximum bound (Myr) | Minimum bound (Myr) |
| --- | --- | --- | --- |
| 1,2,3 | Neognathae / Palaeognathae | 86.5 | 66 |
| 1,2,3,4 | Galloanserae / Neoaves | Free | 66 |
| 1,2,3,4 | Apodidae / Trochilidae | Free | 51 |
| 1,2,3,4 | Sphenisciformes /<br>Threskiornithidae | Free | 60.5 |
| 1,3 | Passerines / Psittaciformes | 65.5 | 53.3 |
| 2,4 | Passerines / Psittaciformes | Free | 53.3 |
| 3 | Oscines / Suboscines | 34 | 28 |
| 4 | Neognathae / Palaeognathae | 140 | 66 |

**Figure S1:**
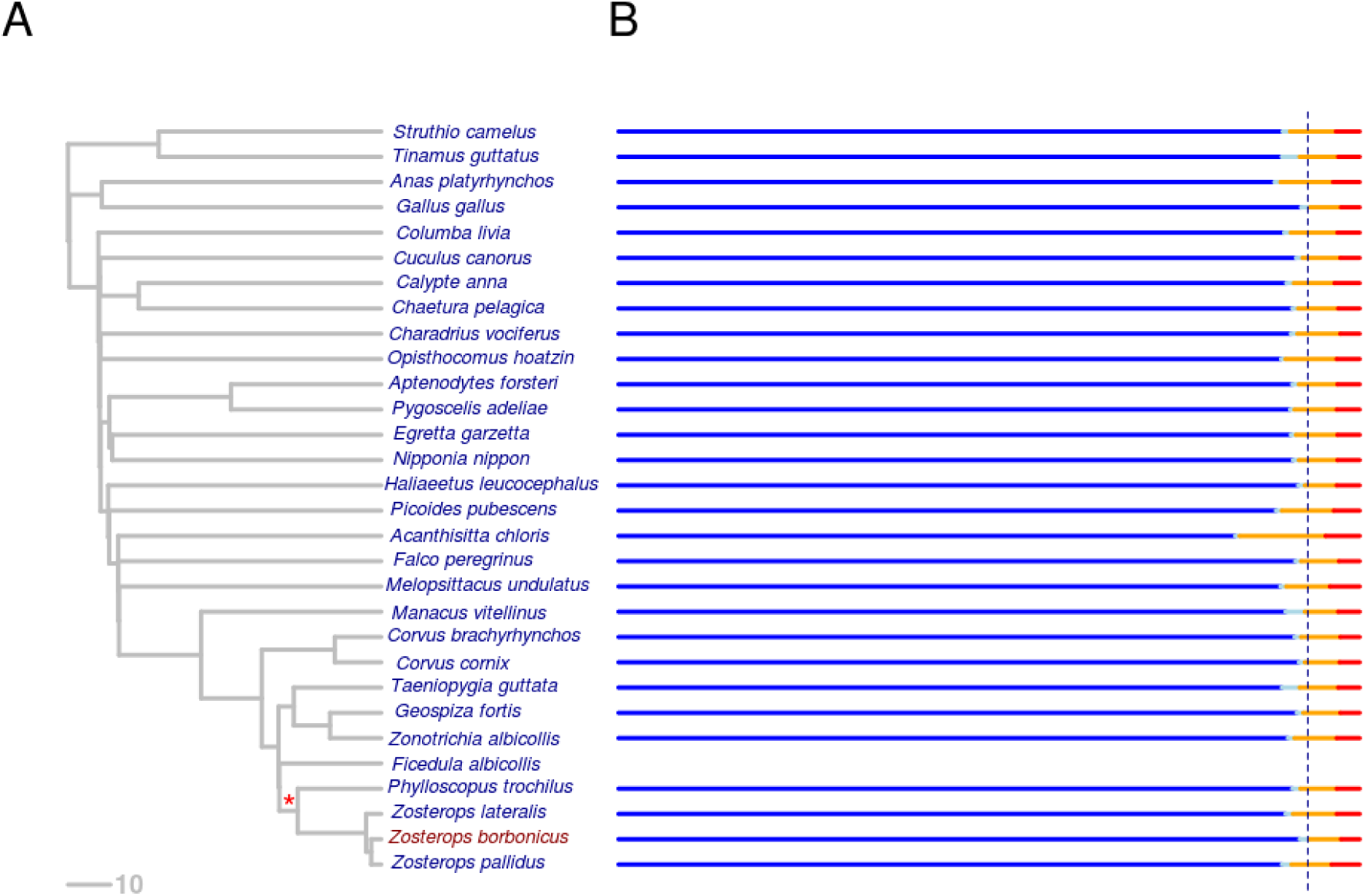
A) Phylogenetic tree based on all investigated avian species. The red star indicates the origin of the two neo-sex chromosomes (ancestor of Sylvioidea). B) Summary statistics of the BUSCO analysis based on the 27 genome assemblies used as input for DeCoSTAR. Blue, light blue, orange and red colors indicate single copy, duplicated, fragmented, missing genes, respectively. The blue dotted line corresponds to the cumulative proportion of complete genes (single copy and duplicated ones) found in *Z. borbonicus*. For unknown reasons, BUSCO analysis failed for the *F. albicollis* assembly.

**Figure S2:**
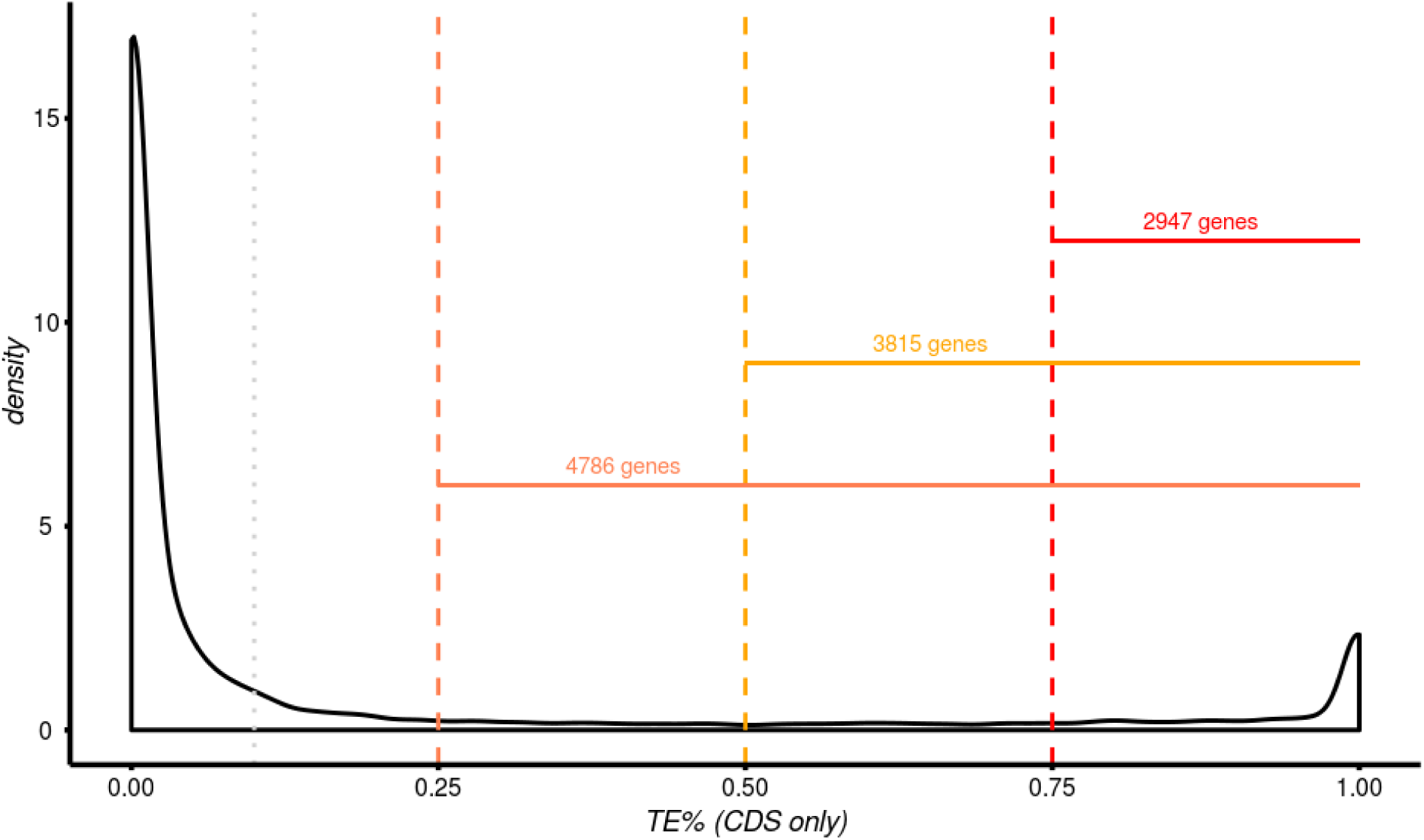
Distribution of the TE content in coding regions observed in the 22,558 *Z. borbonicus* gene models. The grey line shows the threshold used for identifying the most accurate genes (see Fig. 1).

**Figure S3:**
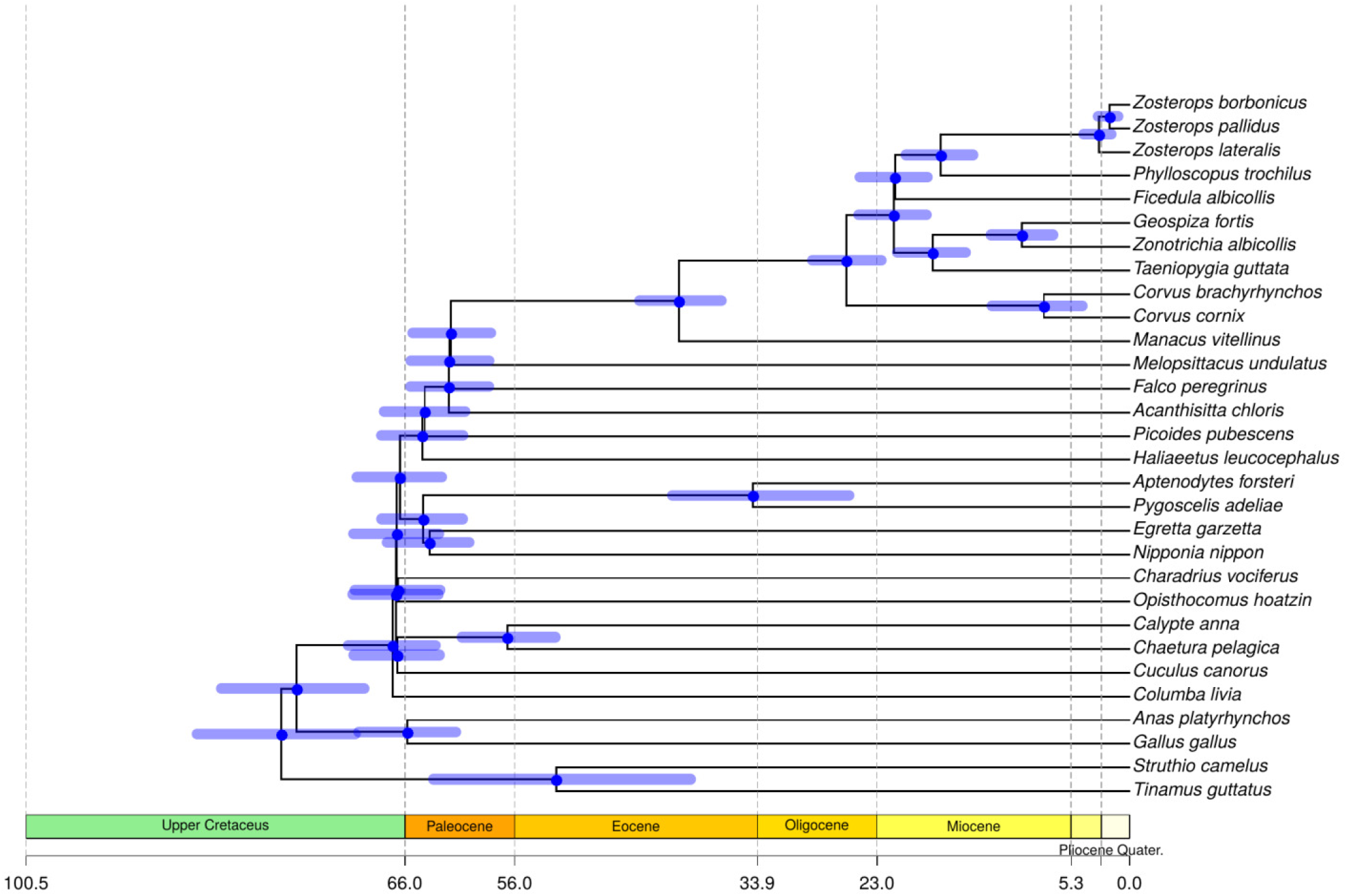
Molecular dating of the 30 birds species. Molecular dating estimates are based on the dataset 1 with CAT GTR substitution model, log-normal molecular rate model and calibration set 1 (Table S3).

**Figure S4:**
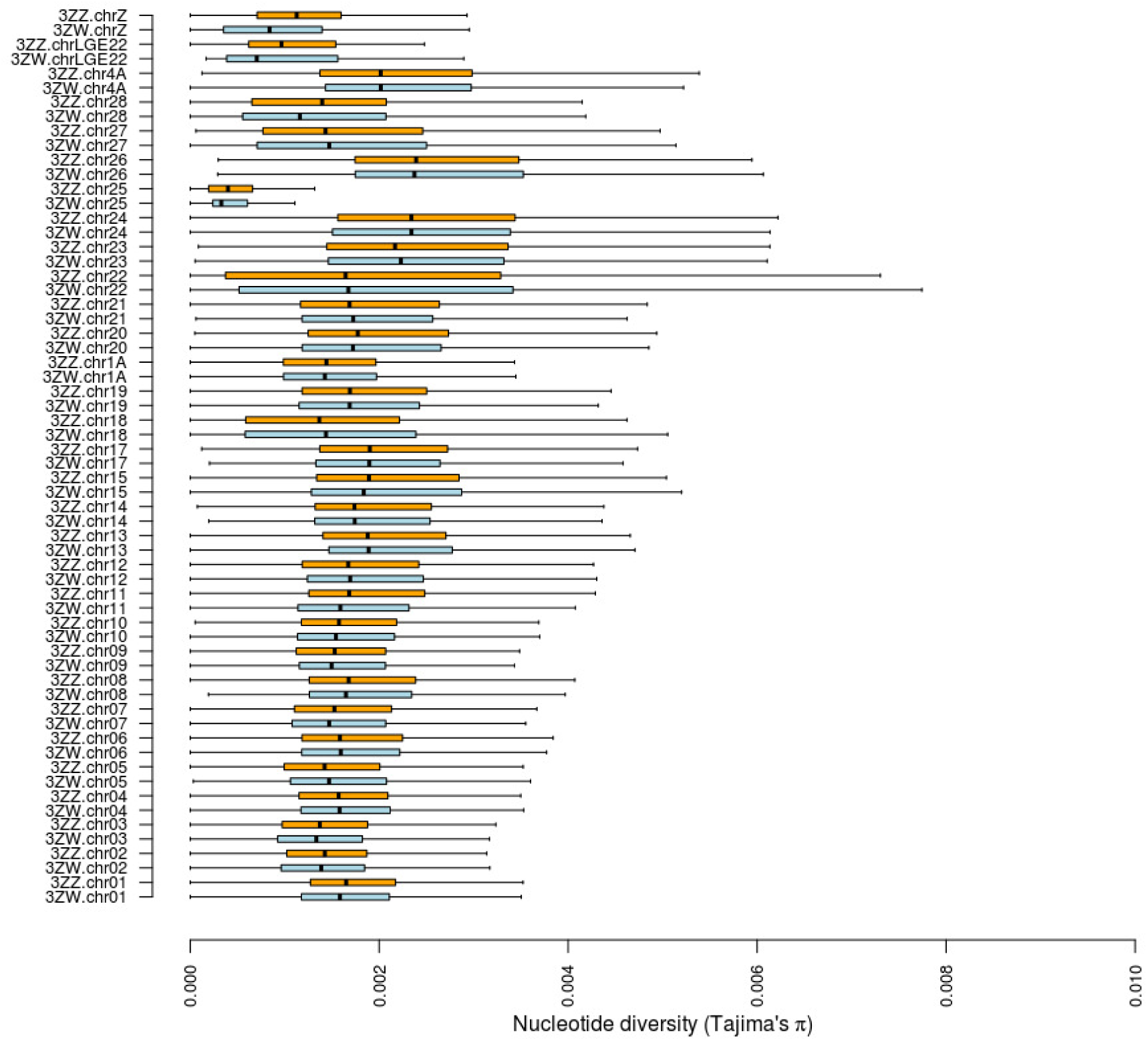
Interchromosomal and interdataset variation in Tajima’s π_females_ (light blue) and π_males_ (orange) over non-overlapping 10-kb sliding windows.

**Figure S5:**
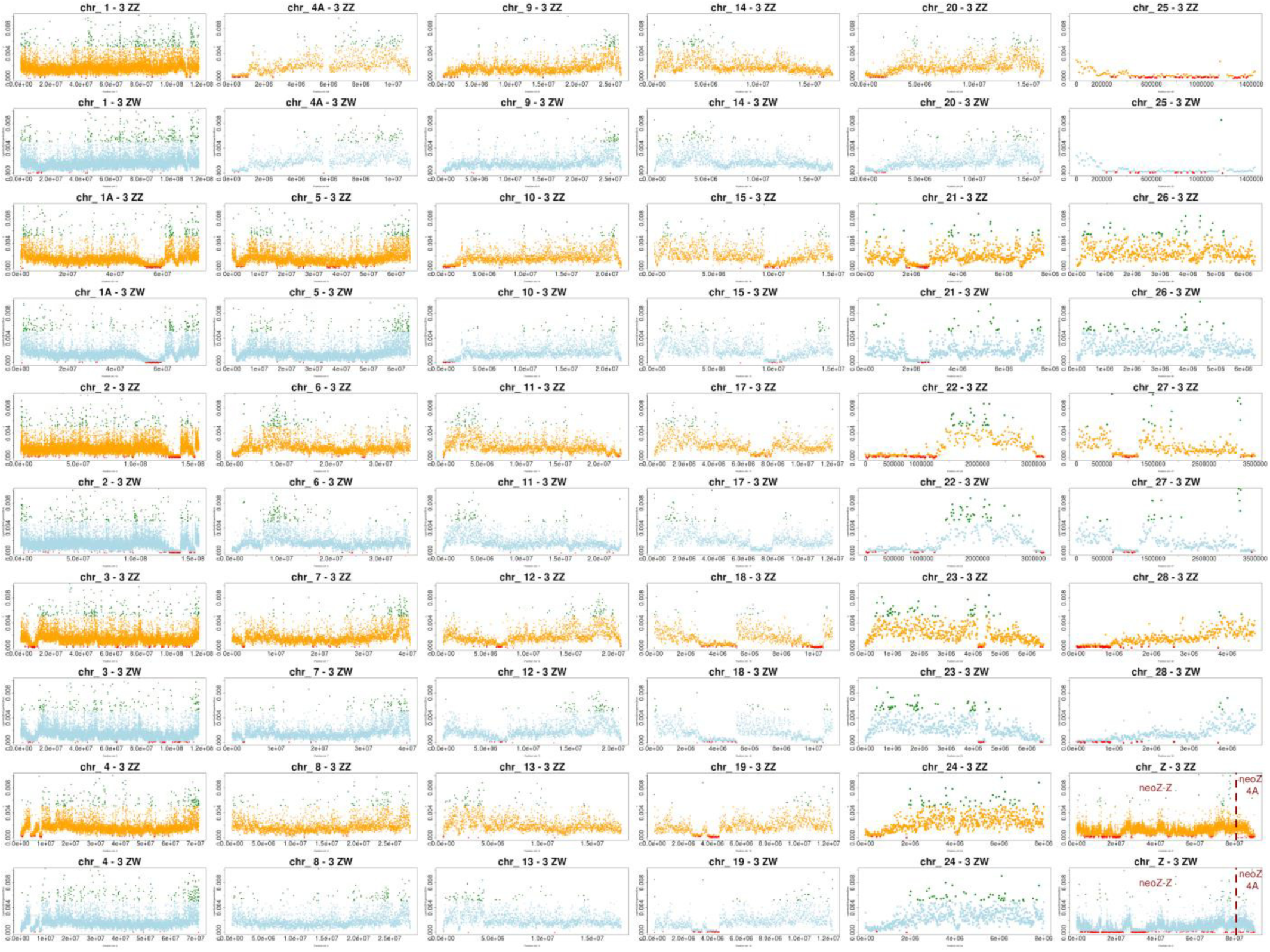
Nucleotide diversity variations along 30 *Z. borbonicus* chromosomes as estimated using genetic information from three males (π_males_, orange) and three females (π_females_, light blue). For each dataset, top 2.5% and bottom 2.5% of π values among windows scanning all chromosomes are shown in green and red, respectively. The chromosomal breakpoint on the neoZ chromosome is shown with a red line.

**Figure S6:**
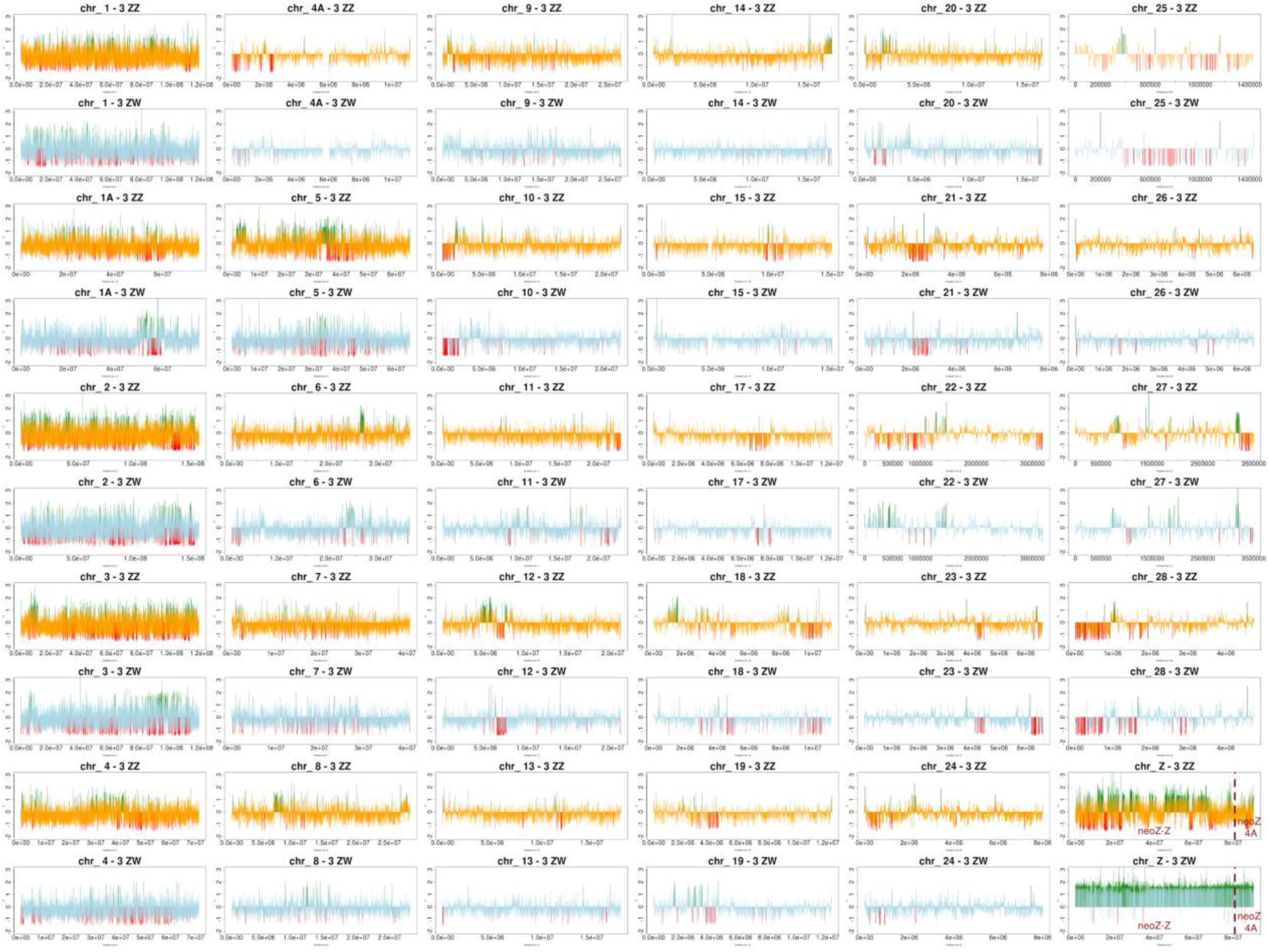
Tajima’s D variations along 30 *Z. borbonicus* chromosomes as estimated using genetic information from three males (orange) and three females (light blue). Top 2.5% and bottom 2.5% of D values among windows scanning all chromosomes are shown in green and red, respectively (baseline for bars is for D=0). The chromosomal breakpoint on the neoZ chromosome is shown with a red line.

**Figure S7:**
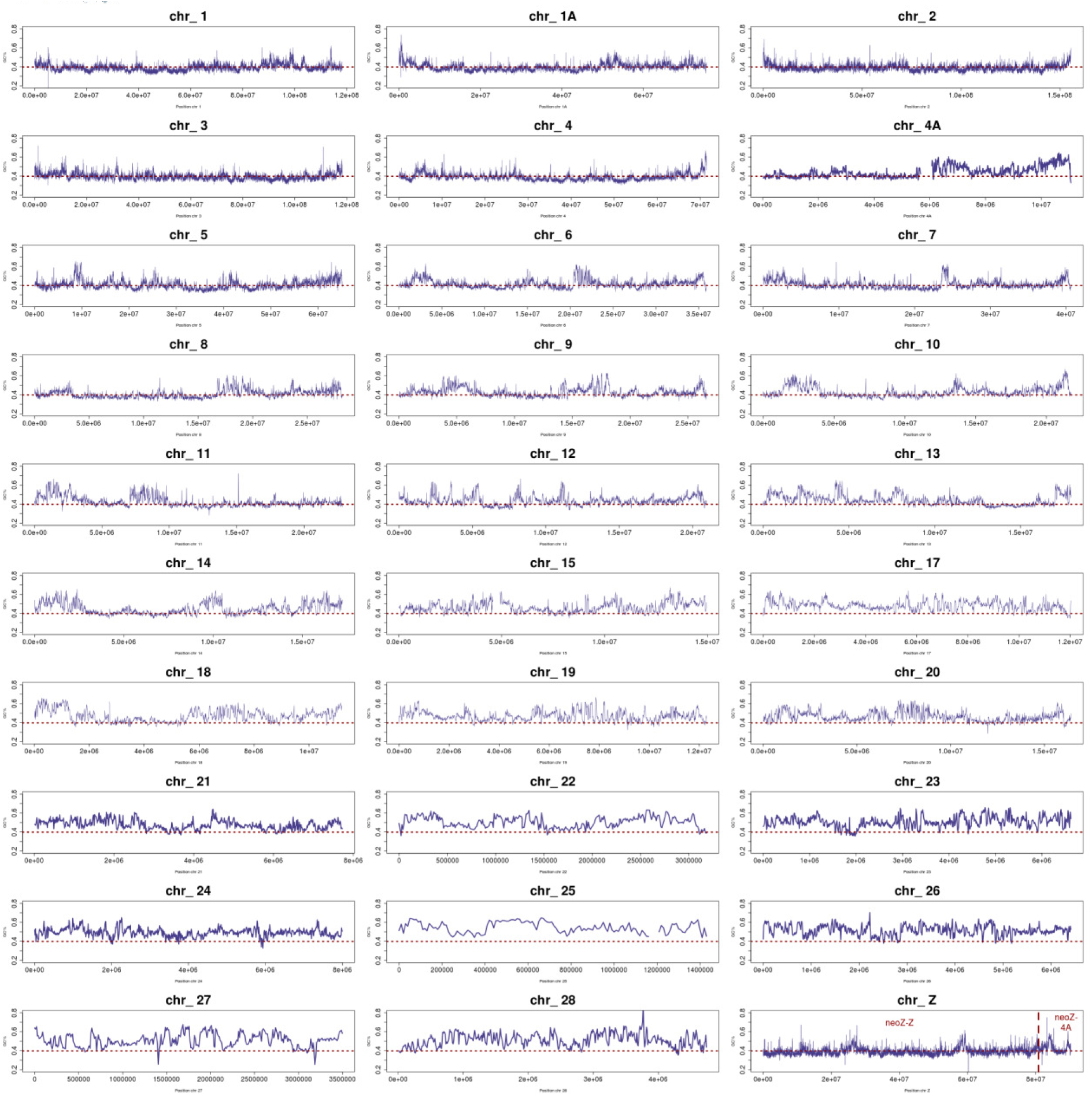
Variation in G+C content along 30 *Z. borbonicus* chromosomes. The red dotted line indicated the median GC value over non-overlapping 10kb sliding windows for scaffolds assigned to chromosomes only. As expected, strong departures from this median values are observed on minichromosomes.

**Figure S8:**
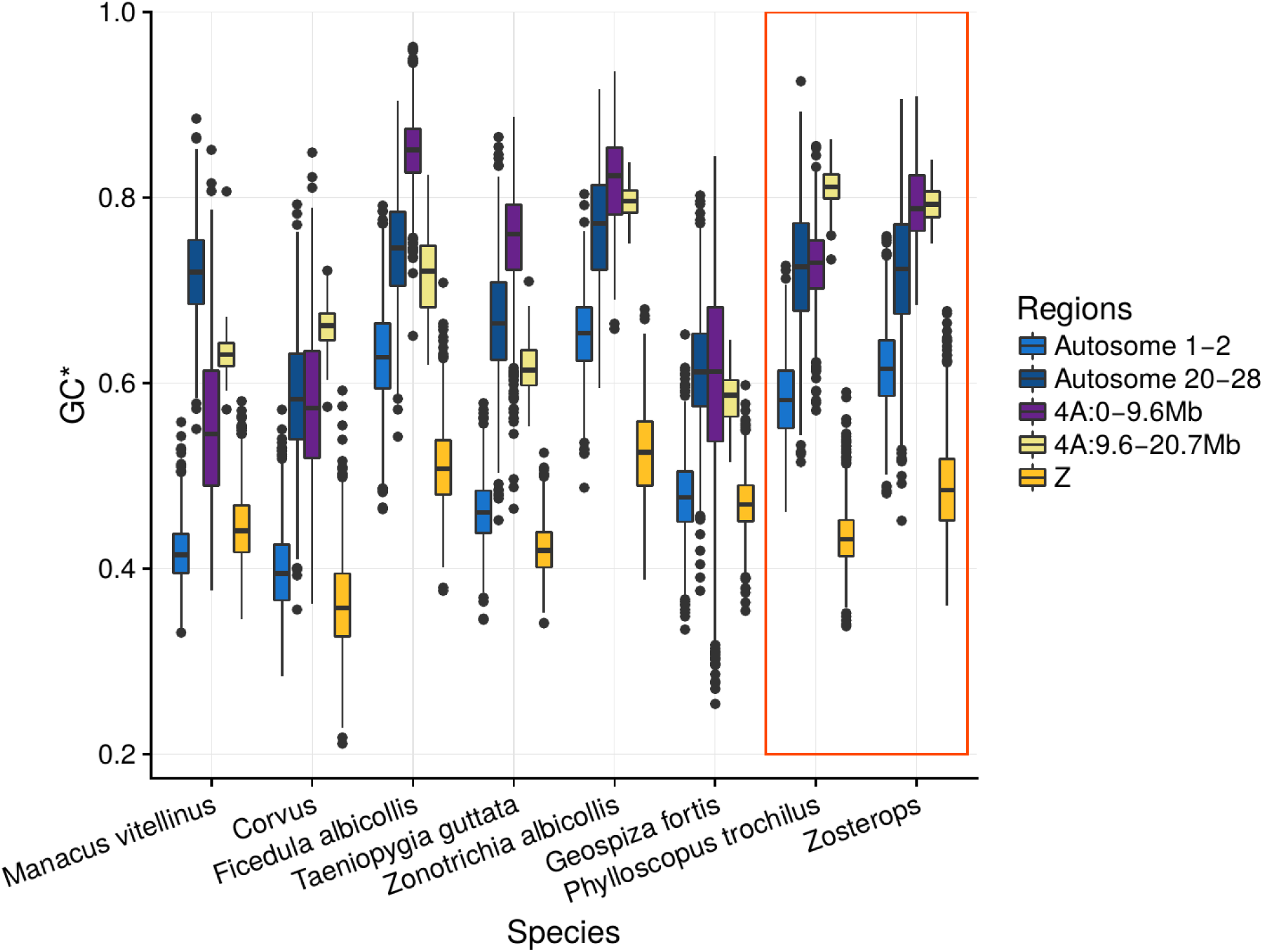
Variations in GC content at equilibrium (GC*) at third codon positions. Variability was obtained by bootstrapping genes within each genomic region.

**Figure S9:**
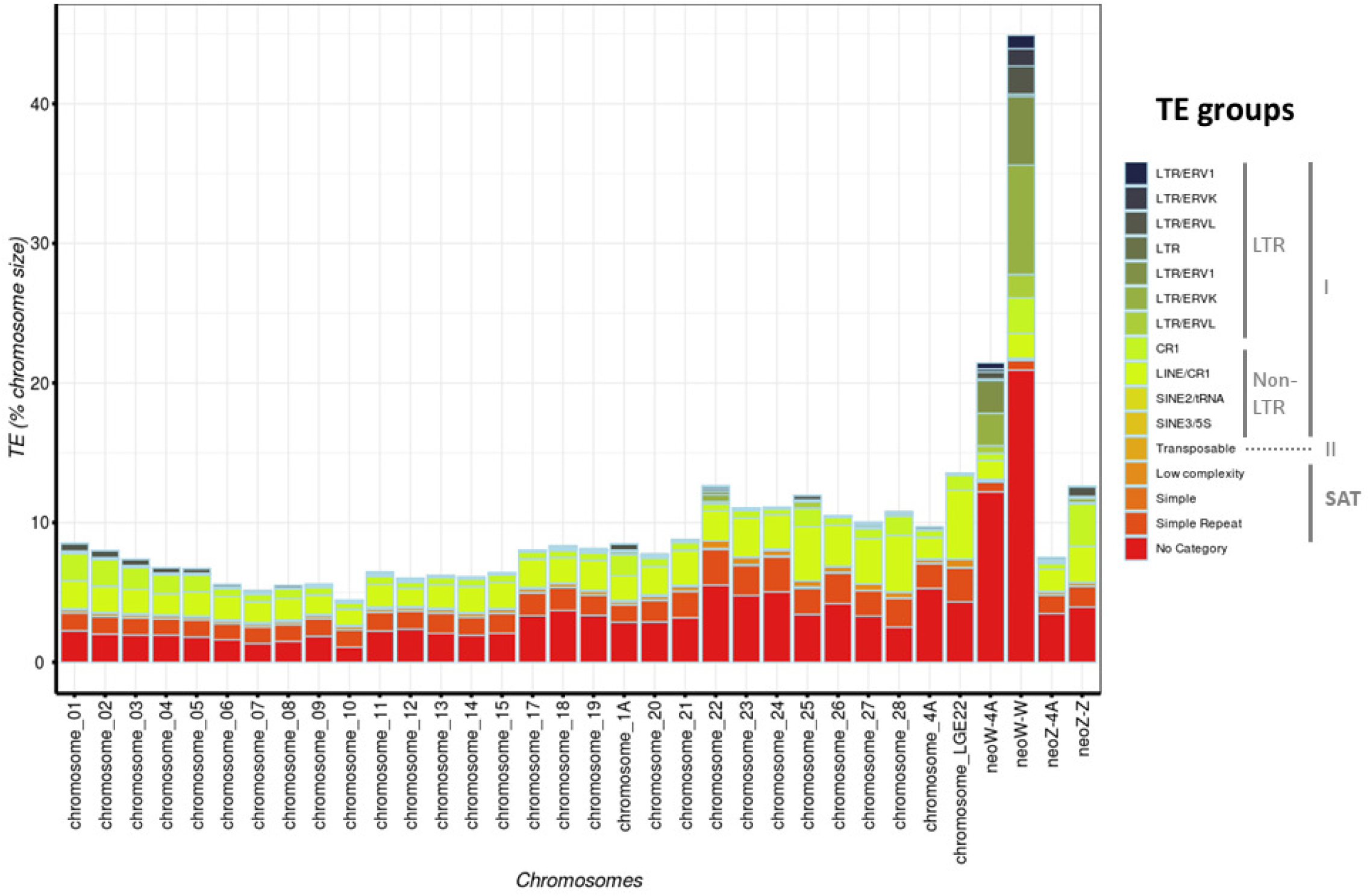
TE density for each chromosome and for each RepBase TE family. Class I TE elements were categorized following the two subclasses: LTR and non-LTR retrotransposons.

